# Tyrosine halogenation converts α-defensins into potent immunomodulators

**DOI:** 10.1101/2023.11.01.565105

**Authors:** Alessandro Foti, Robert Hurwitz, Kathrin Textoris-Taube, Miha Milek, Adam Pickrum, Ashley Castellaw, Hee Jin Kim, Ravishankar Chandrasekaran, Stefan Becker, Venus Singh Mithu, Renate Krueger, Horst von Bernuth, Michael Mülleder, Abin Biswas, Pauline Fahjen, Moritz Winkler, Ulrike Abu Abed, Daniel Humme, Stephanie Thee, Antje Prasse, Benjamin Seeliger, Dieter Beule, Christian Griesinger, Markus Ralser, Victor J. Torres, Arturo Zychlinsky

**Affiliations:** Department of Cellular Microbiology, Max Planck Institute for Infection Biology, Berlin, Germany; Protein Purification Core Facility, Max Planck Institute for Infection Biology, Berlin, Germany; Charité Universitätsmedizin Berlin, Department of Biochemistry, Core Facility - High-Throughput Mass Spectrometry Berlin, Germany; Berlin Institute of Health at Charité–Universitätsmedizin Berlin, Core Unit Bioinformatics, Berlin, Germany; Department of Host-Microbe Interactions, St. Jude Children’s Research Hospital, Memphis, TN, USA; NMR Based Structural Biology, Max Planck Institute for Multidisciplinary Sciences, Göttingen, Germany; Charité - Universitätsmedizin Berlin, corporate member of Freie Universität Berlin, Humboldt-Universität zu Berlin, and Berlin Institute of Health, Department of Pediatric Respiratory Medicine, Immunology and Critical Care Medicine, Berlin, Germany; Labor Berlin Charité-Vivantes, Department of Immunology, Berlin, Germany; Berlin Institute of Health (BIH) at Charité - Universitätsmedizin Berlin; Charité - Universitätsmedizin Berlin, corporate member of Freie Universität Berlin, Humboldt-Universität zu Berlin, and Berlin Institute of Health (BIH), Berlin-Brandenburg Center for Regenerative Therapies (BCRT), Berlin, Germany; Core Facility – High-Throughput Mass Spectrometry, Charité – Universitätsmedizin Berlin, Corporate Member of Freie Universität Berlin and Humboldt-Universität zu Berlin, Core Facility – High-Throughput Mass Spectrometry, Am Charitéplatz 1, Berlin, Germany; Quantitative Biology, Max Planck Institute for Infection Biology, Berlin, Germany; Biological Optomechanics Division, Max Planck Institute for the Science of Light, Erlangen, Germany; Microscopy Core Facility, Max Planck Institute for Infection Biology, Berlin, Germany; Department of Dermatology, Venerology and Allergology, Charité Universitätsmedizin Berlin, corporate member of Freie Universität Berlin, Humboldt-Universität zu Berlin and Berlin Institute of Health, Berlin, Germany; Department of Pediatric Respiratory Medicine, Immunology and Critical Care Medicine, Charité-Universitätsmedizin Berlin, corporate member of Freie Universität Berlin, Humboldt-Universität zu Berlin, and Berlin Institute of Health, Berlin, Germany; Department of Respiratory Medicine, Hannover Medical School, Hannover, Germany; Biomedical Research in End-Stage and Obstructive Lung Disease (BREATH), Hannover Medical School (MHH), German Center for Lung Research (DZL), Hanover, Germany; Charité Universitätsmedizin Berlin, Department of Biochemistry, Berlin, Germany; University of Oxford, The Wellcome Centre for Human Genetics, Nuffield Department of Medicine, Oxford, United Kingdom

## Abstract

Neutrophils are immune cells that eliminate microbes using a powerful arsenal of reactive oxygen species, hypohalous acids, proteases, and antimicrobial peptides — yet how these cytotoxic effectors act in concert remains poorly understood. Here, we identify an unrecognized synergy between oxidative and non-oxidative neutrophil defenses. We show that myeloperoxidase-derived hypohalous acids selectively halogenate α-defensins (HNP1–3) at conserved tyrosine residues. This post-translational modification occurs in both human and rat neutrophils and is prominent in diseases marked by neutrophil-rich inflammation, including cystic fibrosis, bacterial pneumonia, and *Staphylococcus aureus* abscesses. Halogenation increases HNP1 hydrophobicity without altering its structure, reprogramming these peptides into potent immunomodulatory mediators. Using single-cell transcriptomics and a *in vivo* peritonitis model, we demonstrate that halogenated HNP1s amplify immune signaling.

Our analysis of the human haloproteome reveals a novel oxidative mechanism by which hypohalous acids rewire immune protein functions, uncovering an unexpected layer of cooperation between neutrophil effector systems. The discovery of halogenated HNP1s represents, to our knowledge, the first characterization of a protein halogenation that has biologically significance in human immunity.

## Introduction

Neutrophils are the most abundant leukocytes in humans and act as first responders of innate immunity. To execute their function, neutrophils deploy tightly regulated oxidative and non-oxidative antimicrobial mechanisms (*1*,*2*). The oxidative response is driven by the NADPH oxidase (NOX2) complex, which generates superoxide that subsequently dismutates, either spontaneously or via superoxide dismutase 1 (SOD1) (*3*), into hydrogen peroxide (H_2_O_2_). In turn, the heme enzyme myeloperoxidase (MPO) converts H_2_O_2_ into highly reactive hypohalous acids (HOX), such as hypochlorous acid (HOCl) (*4*,*5*). These cytotoxic products are indispensable for pathogen clearance and immune signaling; genetic deficiencies in NOX2 or MPO result in life-threatening infections and immune dysregulation (*4*). In parallel, neutrophils deploy non-oxidative antimicrobial effectors, including serine proteases and antimicrobial peptides (*6*).

Both oxidative and non-oxidative effectors are released into restricted compartments such as phagolysosomes and inflammatory lesions, characterized by high local concentrations of reactive oxygen species (ROS), HOX, peptides and proteins where they likely interact (*7*). While HOX are traditionally viewed as broadly cytotoxic molecules that indiscriminately degrade microbial components, emerging evidence shows that MPO-derived oxidants preferentially react with neutrophil proteins (*8*,*9*). This raises the possibility that hypohalous acids serve not only as microbicidal agents, but also as regulators of neutrophil function through post-translational modification of host molecules.

Here, we demonstrate that MPO catalyzes the selective halogenation of immune proteins under pathophysiological conditions. Specifically, we discovered chlorination and iodination of the α-defensin 1–3 (HNP1–3) at conserved tyrosine residues. HNP1–3 are small, cationic, hydrophobic peptides, which differ only at their N-terminal amino acid (*10*). HNP1-3, originally named “alarmins”, were identified because of their mild antimicrobial and immunomodulatory activity (*11*). Using structural, biophysical, and mass spectrometric analyses, we show that halogenation of HNP1 enhances peptide hydrophobicity without altering its structure. Halogenated HNP1s become potent immunomodulators, induce chemokines and inflammatory responses in immune cells, as shown by cell biological analysis and single-cell transcriptomics. We showed that halogenated HNP1s are proinflammatory in an *in vivo* rat model (*12*), since mice lack these peptides (*13*). Notably, halogenated HNP1-3 are abundant in neutrophil-rich human sputum of patients with cystic fibrosis (CF), bronchoalveolar lavage (BAL) fluid from patients with bacterial pneumonia caused by *Streptococcus pneumoniae* and accumulate within abscesses from patients infected with *Staphylococcus aureus*. Altogether this study reveals a novel immune mechanism where neutrophils harness oxidative halogen chemistry to convert HNP1-3 into strong proinflammatory signals.

## Results

### Neutrophils halogenate HNP1–3

Upon activation, neutrophils generate large amounts of ROS and HOX. Using the HOX-sensitive reporter 3′-(p-aminophenyl)-fluorescein (APF) (*14*), we detected robust HOX production within intracellular phagolysosomes and NETotic vacuoles after stimulation with *S. aureus* or the mitogen Phorbol 12-myristate 13-acetate (PMA), as shown by optical diffraction tomography and confocal microscopy (Fig. 1A). As expected, the inhibitors of NOX2 (DPI) or MPO (4-ABAH) reduced APF fluorescence, confirming that HOX production is driven by ROS (Fig. S1A).

**Figure 1.**
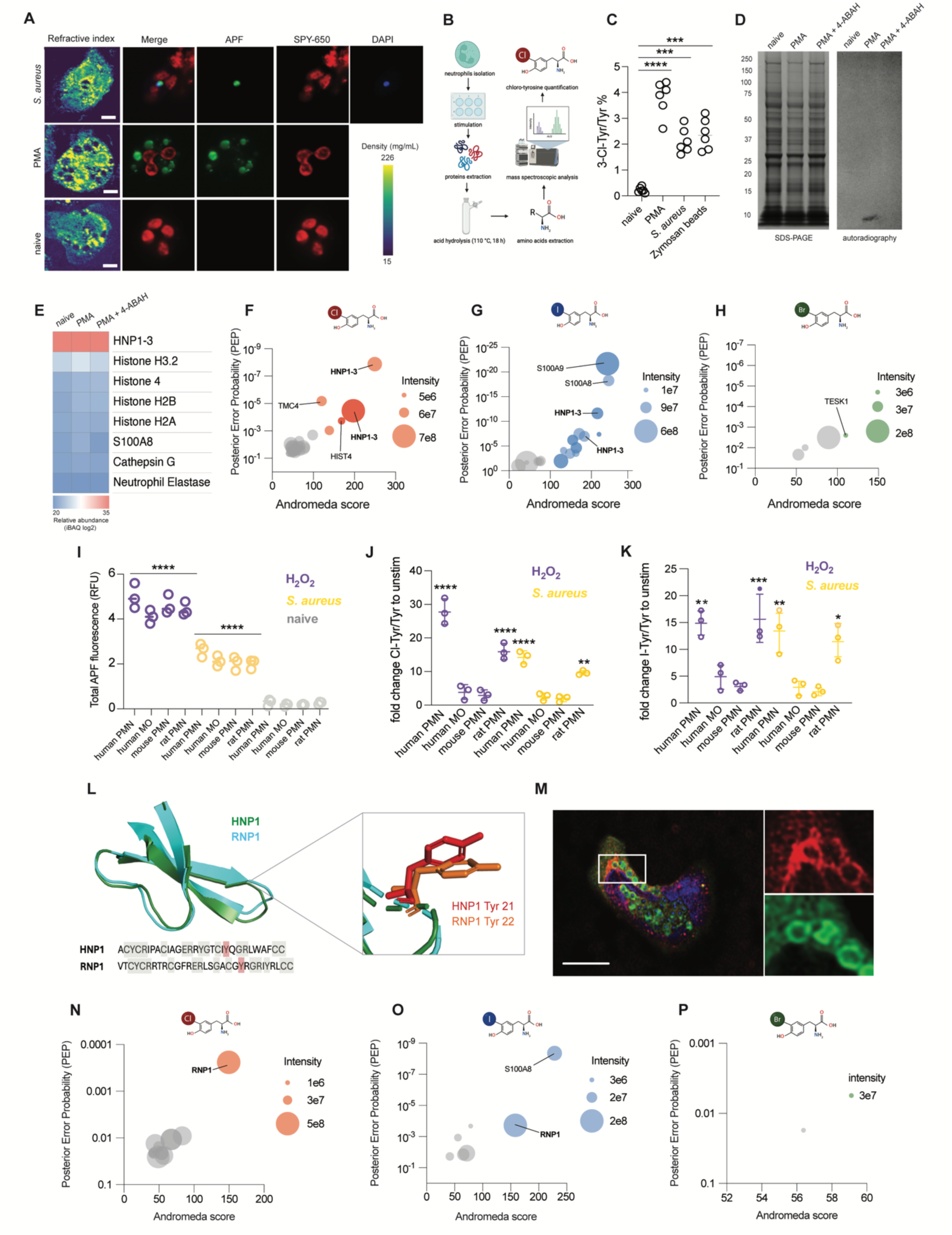
Neutrophils halogenate α-defensin 1–3. (A) Representative optical diffraction tomography (ODT) and confocal microscopy images of human neutrophils activated with PMA (100 nM) or infected with *S. aureus* (MOI=10:1). The density measured by ODT is reported in the color bar on the bottom-right. SPY650 labels DNA and shows the neutrophil lobulated nucleus; APF indicates hypohalous acids; *S. aureus* cells were stained with DAPI before infection. Scale bar, 2 μm. (B) Chloro-tyrosine (ClY) quantification. Human neutrophils were stimulated before protein extraction. The protein extract was hydrolyzed and the amino acids detected by ESI-MS. (C) Ratio of ClY/Y analyzed by hydrolysis and ESI-MS in human neutrophils activated as indicated. Raw data were analyzed by one-way ANOVA with Bonferroni multiple comparisons test; ****p < 0.0001, ***p < 0.001; N=6. (D) SDS-PAGE (left) and autoradiography (right) of ^36^Cl radio-labelled naive and stimulated human neutrophils in the presence or absence of the MPO inhibitor (4-ABAH, 300 µM). Representative experiment of 6 biological replicates (N=6). (E) Relative proteins abundance of proteins in the band (10 kDa) from Fig. 1E analyzed by LC/MS proteomics. (F-H) Mass spectrometry identification of chlorinated (F), iodinated (G) or brominated (H) proteins from human neutrophils stimulated with PMA (dark dots) or *S. aureus* (light dots). The legend indicates the LFQ intensity values for peptide abundance. The chemical structures of chloro-, iodo-, and bromo-tyrosines are shown above their respective panels. Each panel displays a representative experiment of 5 biological replicates (N=5). (I) Detection of hypohalous acids with a APF based assay in human neutrophils and monocytes, as well as in mouse and rat neutrophils, after stimulation with H_2_O_2_ (1 mM) or *S. aureus* (MOI=10:1). Data were analyzed by two-way ANOVA with Tukey’s multiple comparisons test; ****p < 0.0001. N=3. (J-K) Fold change of chloro- or iodo-tyrosines measured by ESI-MS in human neutrophils and monocytes as well as mouse and rat neutrophils stimulated with H2O2 or *S. aureus*, compared to naive cells. Raw data were analyzed by two-way ANOVA with Tukey’s multiple comparisons test, and plotted as fold change. ****p < 0.0001, ***p < 0.001, **p < 0.01, *p < 0.05. N=3. (L) Superimposed 3D-structure of HNP1 (PDB:1gny) and RNP1 (AF-Q62716-F1-model_v2). HNP1 Tyr21 and RNP1 Tyr22 are in very similar positions. The amino acid sequence alignment indicates the conserved residues (gray) between HNP1 and RNP1; red indicates halogenated tyrosine in position 21 in human and 22 in rat. (M) Representative confocal microscopy image of isolated rat neutrophil infected with *S. aureus* and stained for MPO (green), RNP1-3 (red) and DAPI to visualize DNA (blue). Scale bar, 2 μm; Representative image from 3 independent experiments (N=3). (N-P) Mass spectrometry identification of chlorinated (O), iodinated (P) and brominated (Q) proteins from rat neutrophils stimulated with *S. aureus* (MOI=10:1). The legend indicates the LFQ intensity values for peptide abundance in each experiment. Each panel displays a representative experiment of 4 biological replicates (N=4).

To assess protein halogenation, we stimulated human neutrophils with PMA, *S. aureus*, or zymosan beads and quantified 3-chlorotyrosine (ClY)—a stable product of HOCl-mediated protein modification—using acid hydrolysis followed by electrospray ionization mass spectrometry (ESI-MS) (Fig. 1B). ClY accumulated in response to the three stimuli (Fig. 1C). Control experiments showed that the neutrophils lysates did not contain ClY as a byproduct of peptide fragmentation (Fig. S1B), thus strongly suggesting that the modified tyrosine was covalently bound to proteins.

To identify specific halogenated proteins, we labeled neutrophils with radioactive ³⁶Cl, stimulated them, and performed autoradiography after SDS-PAGE separation. Strikingly, a single 10 kDa band was detected (Fig. 1D), which we identified as HNP1–3 by mass spectrometry (Fig. 1E). Fractionation of the neutrophil lysate confirmed that >90% of ClY localized to the fraction enriched in HNP1–3 (Fig. S1C–F). We analyzed the lysate of activated neutrophils by Liquid Chromatography-Tandem Mass Spectrometry (LC-MS/MS) label free quantification (LFQ) and intensity Based Absolute Quantification (iBAQ), using the MaxQuant (*15*) software and searched for halogenated tyrosine containing peptides. This analysis showed that HNP1–3 was chlorinated and iodinated, but not brominated (Fig. 1F–H). Further, Tricine SDS-PAGE combined with LC-MS confirmed chlorination predominantly at Tyr21 and, to a lesser extent, at Tyr16 (Fig. S1G-H). Notably, iodination (iodotyrosine, IY) was favored in Tyr16 (80%) over Tyr21 (20%) (Fig. S1I). Both chlorination and iodination were suppressed by the MPO inhibitor 4-ABAH (Fig. S1J–K). To assess the conservation of HNP1-3 halogenation, we compared human neutrophils, human monocytes, as well as mouse and rat neutrophils. All of these cell types express MPO, however, only human and rat neutrophils express NP1–3 (HNP1–3 in humans and RNP1–3 in rats). Although all MPO-expressing cells produced HOX (Fig. 1I), substantial ClY and IY formation occurred only in human and rat neutrophils, the cell types that express HNP1–3 or their homologs (Fig. 1J–K). The rat orthologs RNP1–3 are similar to human HNP1–3 in both sequence and structure (Fig. 1L). In rat neutrophils, during phagocytosis of *S. aureus*, RNP1–3 and MPO colocalized, indicating their close proximity and consistent with protein halogenation (Fig. 1M). Indeed, proteomic analyses showed that tyrosine 22 in RNP1-2, analogous to HNP1–3 Tyr21, is chlorinated and iodinated, but not brominated (Fig. 1N–P, Fig. S1L). RNP3 halogenation was not detected because the peptide contains a leucine at position 22 (*12*). MS/MS spectra of halogenated HNP1-3 and RNP1-2 peptides corroborated the presence of halogenation on the tyrosine 21 and 16 for chlorination and iodination in humans, respectively, and on tyrosine 22 in rats. Chlorination (+33.9610 Da) or iodination (+125.8966 Da) of tyrosine residues caused a mass shift in the corresponding b- or y-ions compared to the unmodified peptides (Fig. S2).

### The haloproteome of patients reveals HNP1-3 as halogenation target

After identifying HNP1–3 halogenation in healthy donor neutrophils, we investigated this process in neutrophils from patients with defective ROS or HOX production. In chronic granulomatous disease (CGD) (*16*), mutations in NOX2 impair ROS formation(*4*), while in MPO deficiency, ROS are generated but cannot be converted into HOX (*5*). As expected, neutrophils from CGD patients failed to produce ROS upon PMA stimulation, whereas MPO-deficient cells exhibited increased ROS accumulation due to impaired H_2_O_2_ consumption (Fig. 2A). Correspondingly, both patient groups showed minimal HOX production (Fig. 2B). Importantly, reconstitution with exogenous H_2_O_2_ or hypochlorous acid (HOCl) restored halo-tyrosine formation in CGD and MPO-deficient neutrophils, respectively (Fig. 2C), confirming that this post-translational modification requires the ROS–HOX cascade.

**Figure 2.**
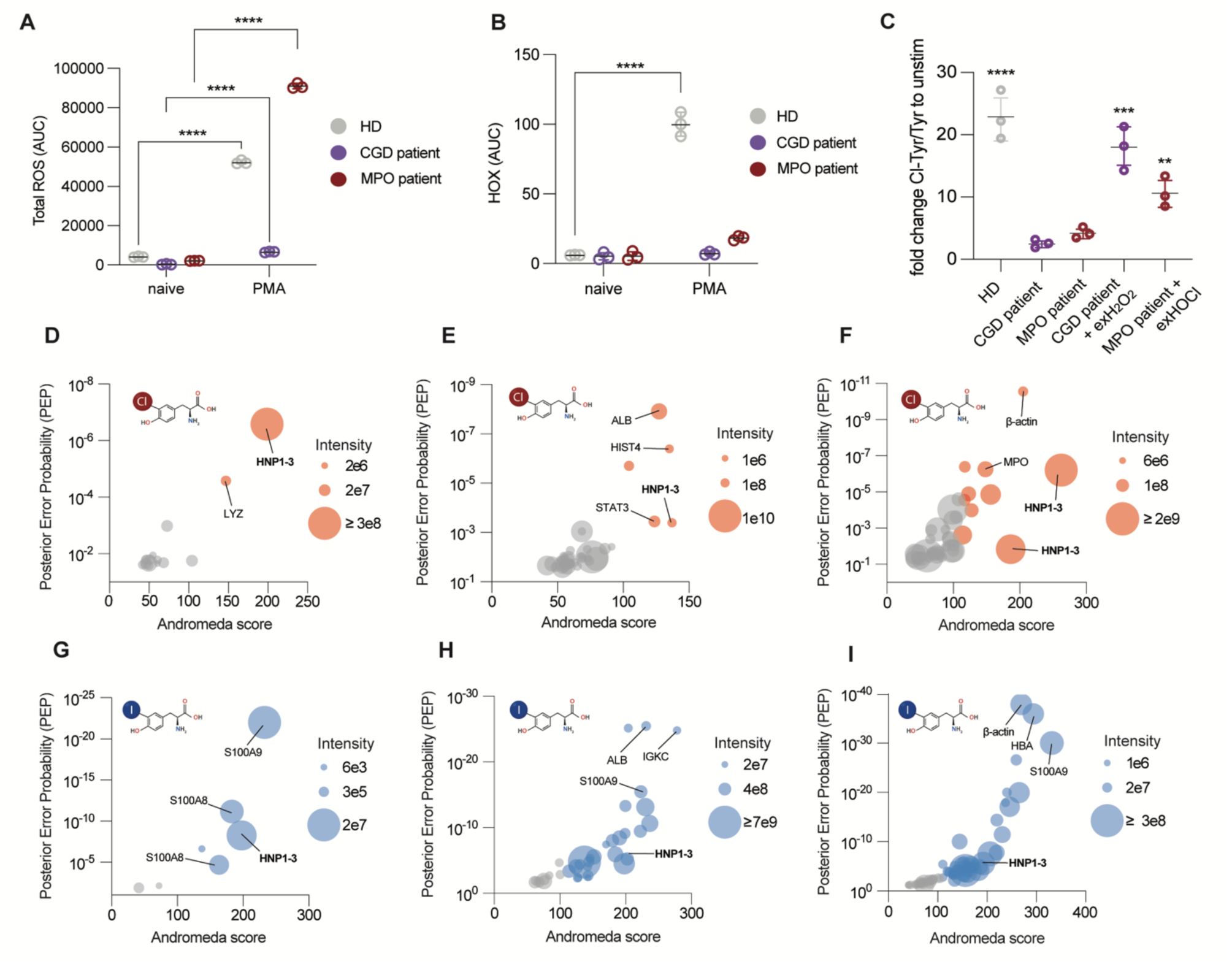
The haloproteome of patients reveals HNP1-3 halogenation. (A) Representative experiment of ROS production, measured with a luminol assay, in naive and PMA (100 nM) stimulated human neutrophils from healthy donors (HD) (N=4), CGD patients (N=3) and MPO deficient patient (N=1) as indicated. The data points in the panel indicate technical replicates. The healthy donor and patient sample assays were performed in parallel. Data were analyzed by two-way ANOVA with Tukey’s multiple comparisons test, and plotted as fold change. ****p < 0.0001. (B) Representative experiment of hypohalous acids formation measured by APF, in naive and PMA simulated neutrophils from HDs (N=4), CGD patients (N=3) and MPO deficient patient (N=1) neutrophils as indicated. The data points in the panel indicate technical replicates. The healthy donor and patient sample assays were performed in parallel. Data were analyzed by two-way ANOVA with Tukey’s multiple comparisons test, and plotted as fold change. ****p < 0.0001. (C) Ratio of ClY/Y analyzed by acid hydrolysis and ESI-MS in HD (N=4), CGD (N=3) or MPO deficient patient (N=1) derived from neutrophils incubated with hydrogen peroxide (1 mM) or hypochlorous acid (500 µM) as indicated. The panel shows a representative of three CGD donors and a single MPO deficient donor. The data points indicate technical replicates. Raw data were analyzed by two-way ANOVA with Tukey’s multiple comparisons test, and plotted as fold change. ****p < 0.0001, ***p < 0.001, **p < 0.01. (D-F) Liquid-Chromatography Mass spectrometry (LC-MS/MS) analysis of (D) cystic fibrosis (CF) patients sputum (N=3), (E) *S. pneumoniae* infected patients BAL (N=4) and (F) *S. aureus* skin infected patients (N=2) for chlorinated proteins. The legend indicates the LFQ intensity values for peptide abundance in each experiment. Peptides with an andromeda score below 100 are shown in grey. (G-I) LC-MS/MS analysis of (G) CF patients sputum (N=3), (H) *S. pneumoniae* infected patients BAL (N=4) and (I) *S. aureus* skin infected patients (N=2) for iodinated proteins. The legend indicates the LFQ intensity values for peptide abundance in each experiment. Peptides with an andromeda score below 100 are shown in grey.

We then assessed tyrosine halogenation of HNP1–3 in clinical samples from patients with neutrophil-dominated infections and inflammation. Using LC-MS/MS, we detected chlorinated Tyr21 (Fig. 2D) and iodinated Tyr16 (Fig. 2G) in HNP1–3 from sputum of cystic fibrosis (CF) patients, which are characterized by chronic neutrophilic airway inflammation (*17*). Additional chlorinated proteins also included lysozyme, histone 4 and transmembrane channel 4, while significantly iodinated targets encompassed the subunits S100A8 and S100A9 of calprotectin, and lysozyme (Fig. S3A). Di-chlorination was also observed on HNP1-3 (Fig. S3A). We then analyzed bronchoalveolar lavage (BAL) fluid of *Streptococcus pneumoniae*-infected patients hospitalized in the intensive care unit. We also observed significant levels of HNP1–3 chlorination at Tyr21 (Fig. 2E) and iodination at Tyr16 (Fig. 2H). Significant iodination was also observed on IGKC (immunoglobulin kappa constant) (Fig. 2H). Additional chlorination was observed on histone H4 and STAT3, with di-chlorination on HNP1–3 Tyr21 and IGL7 Tyr66 (Fig. S3B). Several plasma proteins, including haptoglobin, and serum albumin, also exhibited significant halogenation levels (Fig. S3B). Similar findings emerged in skin abscesses from *S.aureus*-infected patients. Halogenated HNP1–3 were abundant, particularly chlorination at Tyr21 (Fig. 2F) and iodination at Tyr16 (Fig. 2I). Other halogenated proteins were also MPO, β-actin, lactoferrin, and histone H4 (Fig. S3C). Notably, chlorination also targeted Tyr 3 on HNP1 (AC-ClY-CRIPACIAGERR) (Fig. S3C). No halogenated proteins were detected in adjacent non-inflamed tissue (data not shown), confirming a localized phenomenon. Across all three disease contexts, HNP1–3 and Histone H4 consistently emerged as dominant chlorinated targets. Calprotectin subunits S100A8 and S100A9 were notably the most iodinated proteins, highlighting their potential role in modulating inflammation. Bromination of proteins was minimal and did not reach statistical significance.

Gene Ontology (GO) enrichment analysis indicated that the subset of halogenated proteins is functionally enriched in pathways associated with the modulation of inflammation, DNA repair, innate immune defense against microbial pathogens, regulation of adaptive immunity and maintenance of tissue structure and organization (Fig. S3D–F). These associations suggest that halogenation may preferentially occur in proteins that participate in host defense and homeostatic remodeling processes. Altogether, these findings demonstrate that HNP1–3 halogenation occurs robustly in human pathological settings, preferentially targets key immunoregulatory proteins, and likely contributes to shaping inflammatory and antimicrobial responses in disease.

### Halogenation fine-tunes HNP1-3 physicochemistry without structural disruption

Tyrosine halogenation is a stable modification which can alter protein physicochemical properties (*18*). Covalently attached halogens can impact the steric and charge environment around tyrosines, potentially influencing protein interactions. Notably, in HNP1-3, Tyr16 and Tyr21 contribute to the formation of two hydrophobic pockets through interactions with phenylalanine 28 and tryptophan 26 (Fig. S4A). HNP1-3 are short peptides (29–30 amino acids) (*10*) and can be chemically synthesized, allowing for site-specific modifications (*19*). To explore the functional consequences of tyrosine halogenation, we synthesized the halogenated HNP1s which were abundantly found in patient samples, namely the Tyr21 chlorinated HNP1 (ClY21-HNP1) and the Tyr16 iodinated HNP1 (IY16-HNP1), and compared their properties to the native peptide. X-ray crystallography of ClY21-HNP1 revealed that chlorination preserved the overall 3D structure and hydrophobic pocket organization when compared to unmodified HNP1 (Fig. 3A). Aside from a minor displacement at Tyr21 due to the chlorine atom, the positions of other key hydrophobic residues remained mostly unchanged (Fig. 3B–E). Although IY16-HNP1 did not crystallize, NMR spectroscopy confirmed structural conservation, with minimal chemical shift differences and unchanged proximity between halogenated and neighboring residues (Fig. 3F). The majority of observed chemical shift perturbation (CSP) values fall within ±0.08 ppm, which is within the typical range of experimental uncertainty for peak picking. These low perturbation values further support the conclusion that halogenation at Y21 or Y16 does not induce significant structural changes in the HNP1 fold. The proton contact maps for the halogenated Y16 and Y21 are largely similar to those of their unmodified counterparts and Y21 (Fig. S3B-C), indicating that halogenation does not significantly perturb the overall spatial packing of these residues within the HNP1 structure. Nevertheless, some cross-peaks are missing in both halogenated and non-halogenated forms, primarily due to spectral overlap, but possibly also reflecting subtle changes in the local structural environment. These nuances become more apparent when examining the distance correlation plots, particularly for Y16 (Fig. S3B-C), where the cross-peak-derived distances in IY16 deviate from those in ClY21. This divergence may be attributed to alterations in the local hydration network induced by the bulky, hydrophobic iodine atom in the IY16 variant. Such a change could subtly modulate the local structure surrounding Y16. In contrast, the distance correlation for Y21 between ClY21 and I16Y is more consistent, suggesting that chlorination exerts a comparatively minimal effect on the hydration landscape and local conformation.

**Figure 3.**
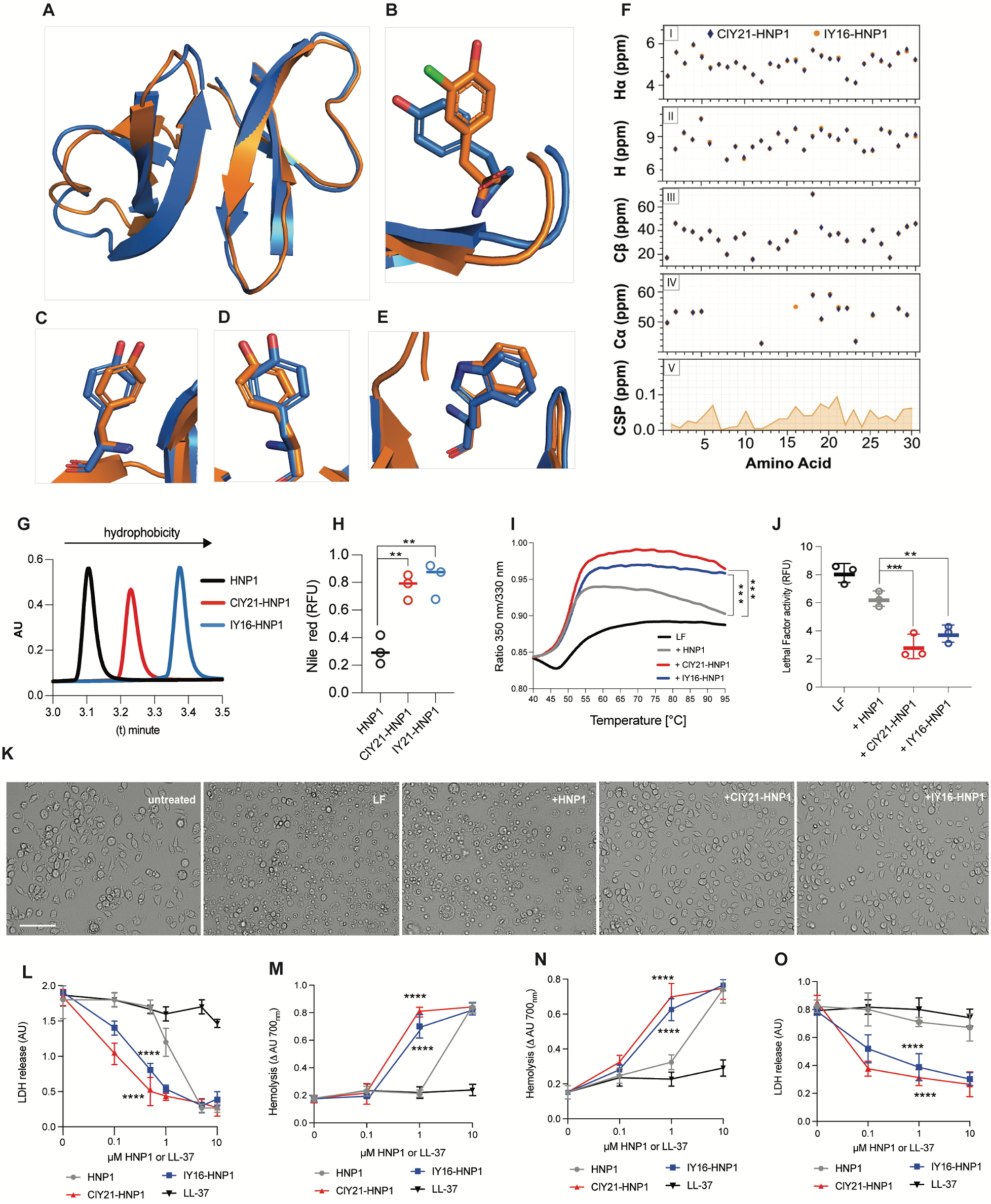
Halogenation fine-tunes HNP1-3 physicochemistry without structural disruption. (A-E) Superimposition of the crystal structure of ClY21-HNP1 (Orange) with HNP1 (Blue) (A) and hydrophobic residues location: Tyr 21 (B), Tyr 16 (C), Tyr 3 (D) and Trp 26 (E). In panel B, chlorine is displayed in green. (F) Panels (I-V) display the comparison of chemical shifts for (I) Hα protons, (II) amide protons, (III) Cβ carbons, and (IV) Cα carbons between ClY21-HNP1 and IY16-HNP1. Chemical shift assignments were derived from two-dimensional 1H–1H TOCSY and 1H–13C HSQC spectra. The structural similarity is further quantified in panel (V), showing the average chemical shift perturbations (CSPs) between ClY21-HNP1 and IY16-HNP1. (G) Representative elution profile of ultra performance liquid chromatography (UPLC) using C18-reverse phase column analysis of HNP1, ClY21-HNP1 and IY16-HNP1; N=3. (H) Fluorescence of HNP1, ClY-21-HNP1 and IY16-HNP1 incubated with Nile red. Data were analyzed by one-way ANOVA with Dunnet multiple comparisons test. N=3. **p < 0.01. (I) Thermostability assay of Lethal Factor (LF) binding to HNP1, ClY-21-HNP1 or IY16-HNP1. Data were analyzed by two-way ANOVA with Tukey’s multiple comparisons test. N=3, ***p < 0.001. (J) Proteolytic measurement of the fluorogenic substrate of LF (20 nM) in presence of HNP1, ClY21-HNP1 or IY16-HNP1 (1µM). Data were analyzed by one-way ANOVA with Dunnet multiple comparisons test. N=3, **p < 0. 01, ***p < 0. 001. (K) Representative images of RAW 264.7 macrophages incubated with LF (20 nM) and with HNP1, ClY-21-HNP1 or IY16-HNP1 (1µM). Scale bar, 50 µm, N=5. (L) Cytotoxicity assay of LF on RAW 264.7 cells measured by LDH release. LL37 serves as a negative control. Data points show means ± SEM. Data were analyzed by two-way ANOVA with Tukey’s multiple comparisons test. N=3, ****p < 0.0001. (M) Red cells hemolysis assay of listeriolysin (LLO) in the presence of HNP1, ClY21-HNP1 or IY16-HNP1. Data points show means ± SEM. Data were analyzed by two-way ANOVA with Tukey’s multiple comparisons test. N=3, ****p < 0.0001. (N) Red cells hemolysis assay of pneumolysin (PLY) in the presence of HNP1, ClY21-HNP1 or IY16-HNP1. Data points show means ± SEM. Data were analyzed by two-way ANOVA with Tukey’s multiple comparisons test. N=3, ****p < 0.0001. (O) Cytotoxicity assay of PVL toxin on human neutrophils in the presence of HNP1, ClY21-HNP1 or IY16-HNP1. Data points show means ± SEM. Data were analyzed by two-way ANOVA with Tukey’s multiple comparisons test. N=3, ****p < 0.0001.

Despite maintaining their overall structure, halogenation increased peptide hydrophobicity. Both ClY21-HNP1 and IY16-HNP1 showed delayed elution in Ultra Performance Liquid Chromatography (UPLC) (Fig. 3G) and higher Nile Red binding and fluorescence compared to unmodified HNP1 (Fig. 3H), as proxy of enhanced hydrophobic properties. Since hydrophobicity is a key determinant of HNP1’s biological activity, particularly its ability to disrupt bacterial toxins by binding hydrophobic regions, we investigated whether halogenation enhanced this function.

Previous work established that HNP1-3 act as anti-chaperones, destabilizing bacterial toxins by engaging their hydrophobic surfaces and promoting unfolding (*20*). Molecular dynamics simulations implicate hydrophobic pockets 1 and 2, organized by Tyr16, Tyr21, Trp26, and Phe28, in this mechanism (*21*) (Fig. S3A). Building on this, we tested halo-HNP1 peptides against several bacterial toxins. Thermal stability assays showed that halogenated HNP1s unfolded *Bacillus anthracis* lethal factor (LF) and *S. aureus* Panton-Valentine Leukocidin (PVL) more effectively than unmodified HNP1 (Fig. 3I, Fig. S5A). Enzymatic assays confirmed that halo-HNP1s inhibited LF proteolytic activity more than the native peptide (Fig. 3J). In functional cell-based assays, halo-HNP1s significantly protected RAW 264.7 macrophages from anthrax lethal toxin-induced cell death (Fig. 3K). LDH release assays quantified this protection, showing a 10-fold increase in activity for chlorinated and an 8-fold increase for iodinated HNP1 relative to the native form (Fig. 3L). Importantly, neither unmodified nor halogenated HNP1s inhibited human neutrophil proteases like elastase, cathepsin G, or MMP-8 (data not shown), suggesting target specificity for bacterial toxins. As expected, LL37, a control cationic peptide without known anti-toxin activity, had no effect (Fig. 3L). We extended these findings to other pore-forming toxins, including *Listeria monocytogenes* listeriolysin O (LLO), *S. pneumoniae* pneumolysin (PLY), and *S. aureus* PVL. Erythrocyte lysis assays showed that both LLO and PLY were more susceptible to halo-HNP1s than to unmodified defensin (Fig. 3M–N). Similarly, halogenated peptides were more effective in reducing PVL-mediated human neutrophil toxicity (Fig. 3O). Finally, we tested halo-HNP1 activity against additional staphylococcal toxins (HlgAB, HlgCB, LukAB CC45) and *Clostridioides difficile* toxin B (TcdB). Halo-HNP1s neutralized HlgAB and HlgCB but showed no activity against LukAB CC45 or TcdB (Fig. S5B–E). These results highlight that halogenation subtly reshapes the physicochemical properties of HNP1-3 without compromising their structure, enhancing their antitoxin potential.

### Halogenation enhances HNP1 immunomodulation

HNP1-3 were originally identified for their antimicrobial properties in hypotonic solutions, where they exhibit bactericidal effects (*22*,*23*). To assess whether halogenation impacts killing activity, we tested the bactericidal and bacteriostatic properties of halo-HNP1s against *S. aureus*, *Escherichia coli*, and *C. albicans*. Halogenation did not alter the killing efficiency in hypotonic media (Fig. S6A-C), nor did it affect growth inhibition in nutrient-rich media (Fig. S6D-F). We next examined whether halo-HNP1s act as priming factors for neutrophils. Pre-incubation of human neutrophils with HNP1, ClY21-HNP1, IY16-HNP1, or granulocyte-monocyte colony stimulating factor (GM-CSF) as a positive control, followed by challenge with *S. aureus*, revealed priming effects for both modified and unmodified peptides, with no significant differences among them (Fig. S6G).

HNP1-3 are mild chemotactic agents for immune cells in isotonic, physiologic conditions (*22*,*24*). Using transwell migration assays, we found that halo-HNP1s were significantly more potent chemoattractants than the native peptide for both neutrophils and monocytes (Fig. 4A-B). Notably, IY16-HNP1 increased neutrophil migration sevenfold and monocyte migration two to threefold compared to unmodified HNP1 (Fig. 4A). Cl21Y-HNP1 increased neutrophil migration fivefold and monocyte migration twofold compared to unmodified HNP1 (Fig. 4B). This chemotactic activity was dependent on the intact folded structure, as halo-HNP1s reduction with DTT abrogated migration. Since HNP1-induced chemotaxis is mediated via G protein-coupled receptors (GPCRs) (*25*,*26*) we assessed downstream signaling by measuring ERK1/2 phosphorylation. Both ClY21-HNP1 and IY16-HNP1 induced robust ERK1/2 activation in monocytes, compared to HNP1 (Fig. 4C). This response was abolished by pertussis toxin, confirming GPCRs involvement (Fig. 4C). Next, we explored the functional impact of halo-HNP1s on human monocyte-derived macrophage (HMDM) activity. Previous studies showed that exposure to HNP1-3 enhances the phagocytic capacity of human monocyte-derived macrophages (HMDMs) (*27*). To asses this, we used a *C. albicans* strain constitutively expressing red fluorescent protein (RFP) under the promotor of *Eno1* (*C. albicans*^RFP+^), coincubated with HMDM in presence or absence of HNP1s, and quantified phagocytosed yeast cells. Treatment of HMDMs with halo-HNP1s significantly enhanced their phagocytic uptake of *C. albicans*^RFP+^, doubling the effect seen with unmodified HNP1 (Fig. 4D-E, Fig. S6H).

**Figure 4.**
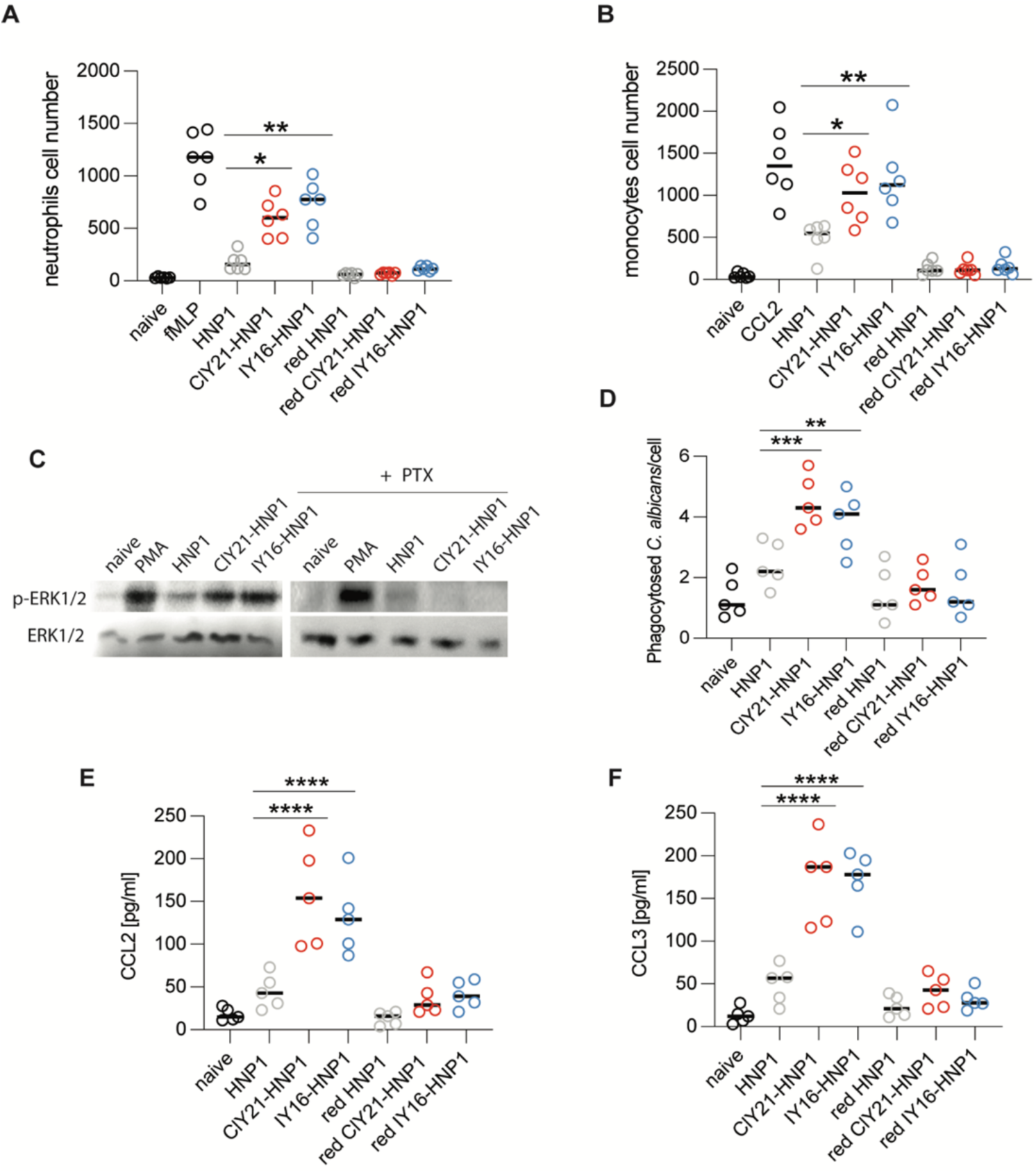
Halogenation enhances the immunomodulatory activity of HNP1. (A-B) Chemotaxis assay of human neutrophils (A) or monocytes (B) stimulated with HNP1, ClY21-HNP1 or IY16-HNP1. Red HNP1, red ClY21-HNP1 and red IY16-HNP1indicates peptides reduced with DTT and serve as negative controls. fMLP and CCL2 are positive chemotaxsis controls for neutrophils and monocytes respectively. Data were analyzed by two-way ANOVA with Tukey’s multiple comparisons test; N=6; ***p < 0.001, **p < 0.01, *p < 0.05. (C) Western blot analysis of phospho-ERK1/2 and ERK1/2 levels in human monocytes incubated with HNP1, ClY21-HNP1 or IY16-HNP1 for 2 h (N=3). PMA was used as a positive control. PTX indicates monocytes pre-treated with Pertussis toxin. (D) Representative quantification of phagocytosed *C. albicans* by HMDM after 4 hours post-infection. Data were analyzed by two-way ANOVA with Tukey’s multiple comparisons test; N=3; ***p < 0.001, **p < 0.01. (E-F) CCL2 (E) and CCL3 (F) extracellular concentration released from HMDM stimulated with HNP1, ClY-21-HNP1 and IY16-HNP1 for 4 h. Red HNP1, red ClY21-HNP1 and red IY16-HNP1indicates peptides reduced with DTT and serve as negative controls. Data were analyzed by one-way ANOVA with Tukey’s multiple comparisons test; N=5; ****p < 0.0001.

To assess whether halo-HNP1s induce cytokine secretion, HMDMs were treated with HNP1 and halo-HNP1s for 4 hours, and chemokine levels in the supernatant were subsequently quantified by ELISA. The analysis revealed a statistically significant increase in the secretion of CCL2 (MCP-1) and CCL3 (MIP-1α) compared with cells exposed to the unmodified HNP1 peptide (Fig. 4E-F). These findings indicate that halogenation enhances the immunomodulatory properties of HNP1, augmenting its ability to activate macrophages and promote the release of chemotactic mediators that recruit monocytes and other immune cells.

### Halo-defensins are potent proinflammatory agents

To explore the immunomodulatory potential of halogenated defensins on circulating immune cells, we performed single-cell RNA sequencing (scRNA-seq) on peripheral blood mononuclear cells (PBMCs) obtained from four healthy donors. Cells were treated for 4 hours with vehicle, HNP1, ClY21-HNP1, or IY16-HNP1 to capture early transcriptional responses. Principal component analysis (PCA) identified one donor as an outlier, likely due to technical issues during processing, which was subsequently excluded from downstream analyses (Fig. S7A). We then analyzed the three donors following standard scRNA-seq data processing, which included reference mapping to the human PBMC atlas. Cells were annotated at two levels of granularity, enabling the identification of major leukocyte subsets and showed cell proportions that were consistent with expectations (Fig. 5A). Quality control metrics including reference mapping prediction scores, cell-type proportions, marker gene expression and PCA analysis of aggregated read counts confirmed high data quality and accurate cell-type annotation (Fig. S7B–E).

**Figure 5.**
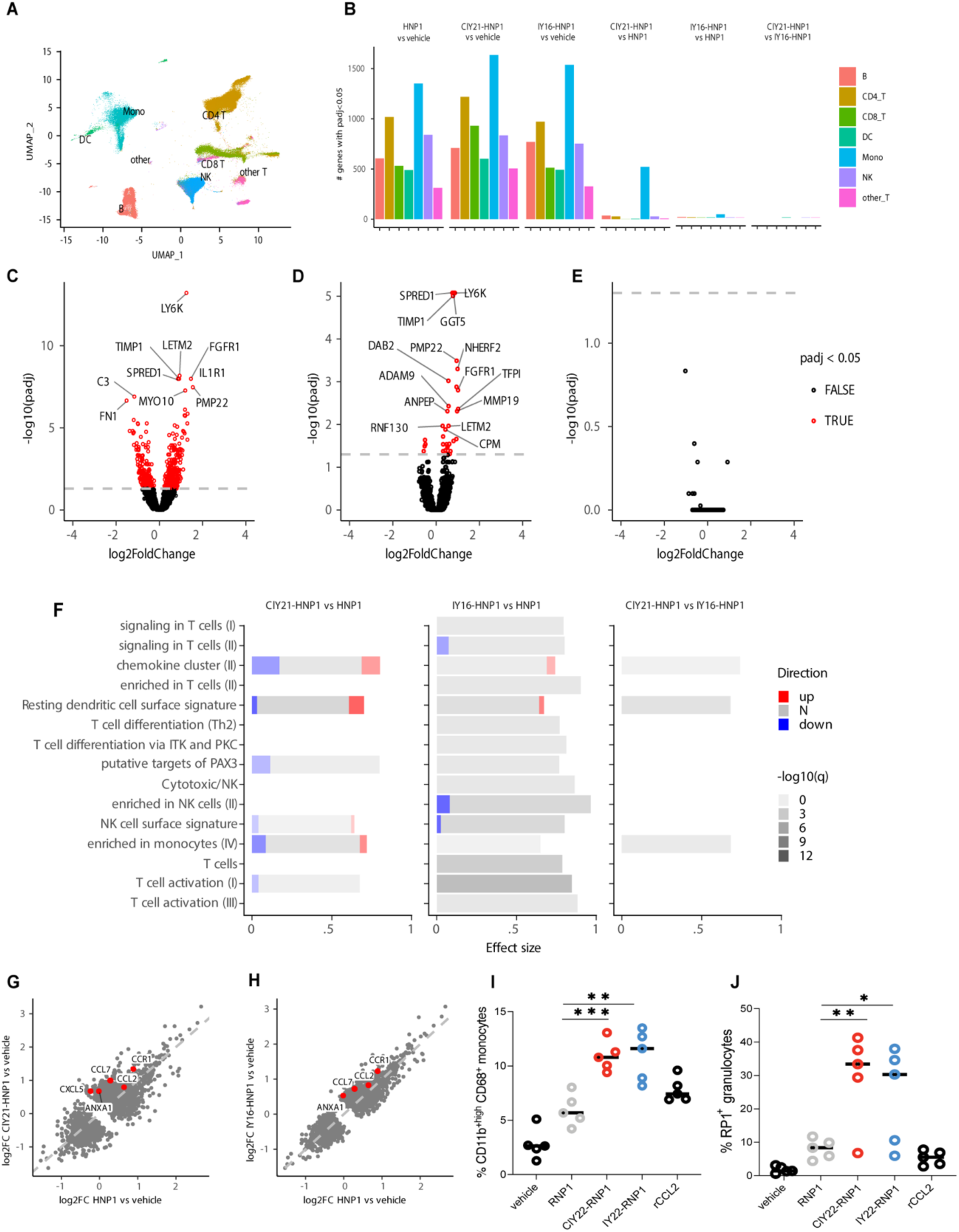
Halo-HNP1s are potent proinflammatory agents. (A) Single cell transcriptomic analysis of human peripheral blood mononuclear cells (PBMCs), incubated with HNP1, ClY21-HNP1 or IY16-HNP1. UMAP embedding of cell types detected using reference mapping from the human PBMC atlas. Cell clusters are indicated in different colors. The analysis includes a total of 200741 cells obtained from 3 healthy donors (N=3). (B) The bar plot displays the count of significantly differentially expressed genes (Wald test, adjusted P value < 0.05) in B cells (B), CD4 T cells (CD4_T), CD8 T cells (CD8_T), Dendritic Cells (DC), Monocytes (Mono), Natural Killer cells (NK), and other T cells (other_T). The color of each bar indicates the specific cell type, as detailed in the legend. (C-E) Log2-transformed fold changes from the pseudobulk differential expression analyses for ClY21-HNP1 versus HNP1 (C), IY16-HNP1 versus HNP1 (D) and ClY21-HNP1 versus IY16-HNP1 (E) in monocytes plotted versus negative log10-transformed adjusted P values (Wald test). Differentially expressed genes (adjusted P value < 0.05) are marked in red, while the non-differential (adjusted P value > 0.05) are shown in black. (F) Gene set enrichment analysis of differentially expressed genes in monocytes for the indicated comparisons (ClY21-HNP1 vs. HNP1, IY16-HNP1 vs. HNP1 and ClY21-HNP1 vs. IY16-HNP1). Bars represent relative fractions within each of the gene sets for mRNAs with decreased (blue) expression, non-differential (N) genes and increased (red) expression (determined by the adj. P value threshold of 0.05). Adjusted P values (q) for the gene set enrichment analyses correspond to the intensity of the bars, i.e. the lower the P value, the higher the bar intensity. Gene sets with AUC > 0.70 and q < 0.0005 are shown. (G-H) Scatter plots of log2-transformed fold changes (log2FC) from pseudobulk differential expression analyses in monocytes. Dots represent all differentially expressed genes (Wald test, adj. P value < 0.05) from the ClY21-HNP1 vs. vehicle (G) and IY16-HNP1 vs. vehicle (H) comparisons. Genes indicated by text labels and red dots belong to the “chemokine cluster (II)”. (I-J) Cell counts expressed in percentage of peritoneal infiltrated monocytes (CD11bhigh/CD68^+^) (I) or granulocytes (RP1^+^) (J) 24 hours after peritoneal injection of RNP1, ClY22-RNP1, IY22-RNP1 or recombinant rCCL2 (as positive control) in rats. Cells were analyzed by flow cytometry. Statistical significance was analyzed by one-way ANOVA with Tukey’s multiple comparisons test, and plotted as percentage. N=5. ***p < 0.001, **p < 0.01, *p < 0.05.

Differential expression analysis revealed robust changes in gene expression upon treatment with HNP1 and its halogenated derivatives. Gene set enrichment analysis demonstrated that both ClY21-HNP1 and IY16-HNP1, as well as the unmodified HNP1, strongly activated innate immune and antiviral transcriptional programs across multiple leukocyte subsets (Fig. 5B, Fig. S8A–C). Pathways related to interferon signaling, type I interferon responses, and general antiviral defense were consistently upregulated in both myeloid (monocytes, dendritic cells) and lymphoid (NK, B, and T cells) cell compartments. Specifically, 1,350, 1,635, and 1,537 differentially expressed genes (DEGs) were regulated by HNP1, ClY21-HNP1, and IY16-HNP1 versus vehicle, respectively, with 742 DEGs shared among all three peptides treatments, which may represent a conserved HNP1 transcriptional response signature. Pairwise comparisons revealed the largest overlap between ClY21-HNP1 and IY16-HNP1 (1,177 DEGs), followed by HNP1 versus IY16-HNP1 (928) and HNP1 versus ClY21-HNP1 (781), suggesting that halogenated HNP1s share extensive similarity while amplifying HNP1’s activity (Fig. S9A).

Monocytes emerged as the most responsive immune cell population when comparing halogenated HNP1s to vehicle or the unmodified peptide (Fig. S9B, Fig. 5B). ClY21-HNP1 treatment resulted in the highest number of differentially expressed genes in monocytes (Fig. 5C), followed by IY16-HNP1, which elicited a less pronounced effect (Fig. 5D). The comparison between ClY21-HNP1 and IY16-HNP1 showed no significant differences (Fig. 5E). Both halogenated peptides promoted the upregulation of key antiviral and proinflammatory genes, including LY6K, FGFR1, and IL1R1, while downregulating immunoregulatory factors such as TIMP1 and C3, indicating a substantial remodeling of receptor signaling and immune effector pathways.

Gene set enrichment analysis further highlighted functional consequences of halogenation (Fig. 5F). The ClY21-HNP1 vs HNP1 comparison resulted in the detection of DEGs which were enriched for pathways related to chemokine production, dendritic cell surface signatures, and T-cell and monocyte activation. The IY16-HNP1 vs HNP1 comparison showed similar trends, with minor differences, including slight downregulation of NK cell–associated signatures. Scatter plot visualization of DEGs revealed increased level of several chemokine cluster genes, including CCL2, CCL7, CXCL5, and CCR1, compared to HNP1 treatment alone (Fig. 5G–H). Notably, CCR1 was also upregulated in CD4⁺ T cells following halogenated HNP1s treatment (Fig. S10), suggesting broader enhancement of chemotactic signaling beyond the myeloid compartment. Together, the differential gene expression analyses in monocytes show that all three peptides elicit a robust interferon-associated transcriptional response, with the halogenated HNP1 variants further driving a distinct and more potent innate immune response compared to the unmodified peptide. Next, we evaluated whether these immune-stimulating effects translated into functional immune cell recruitment *in vivo* in a peritonitis model in Sprague-Dawley rats. Using an intraperitoneal inflammation model to test leukocyte migration, we compared the chemotactic activity by injecting vehicle, unmodified RNP1, ClY22-RNP1, IY22-RNP1, and recombinant rat CCL2 (rCCL2) as a positive control (Fig. 5I). After 24 hours post-injection, the intraperitoneal infiltrated cells were isolated. Flow cytometric analysis of recruited immune cells revealed that both ClY22-RNP1 and IY22-RNP1 induced significantly higher granulocyte and CD11b^high^ CD68^+^ monocyte infiltration in rats compared to vehicle and unmodified RNP1 (Fig. 5J-K). Notably, halogenated RNP1 displayed chemotactic potency higher than rCCL2, a classical innate immune cell chemoattractant (Fig. 5J-K). These data demonstrate the potent immunomodulatory function halogenated HNP1s *in vivo*.

## Discussion

Neutrophils are indispensable effector cells of the innate immune system, equipped with a broad repertoire of antimicrobial mechanisms that encompass both oxidative and non-oxidative strategies. Traditionally regarded as terminal microbial killers, neutrophils are now recognized as sophisticated immunomodulators capable of shaping local and systemic immune responses through the release of cytokines, oxidants and antimicrobial proteins. Yet, how these distinct effector arms interact within the confined and chemically reactive microenvironments of phagolysosomes, inflammatory lesions, or extracellular traps has remained largely unresolved (*28*). In this study, we uncover a cooperative mechanism linking neutrophil oxidative activity to the functional diversification of immune proteins. We show that oxidative halogenation, mediated by MPO-derived hypohalous acids (HOX), chemically transforms α-defensins into potent immunoregulatory mediators, revealing a novel intersection between oxidative halogen chemistry and innate immune signaling.

Halogens are highly reactive elements with diverse and profound biological roles. Chloride is essential for cellular osmotic balance and gastric acid production, fluorine contributes to bone and dental health, and iodide is critical for thyroid hormone synthesis. In contrast, the biological functions of bromide remain poorly characterized (*29*,*30*,*31*). Within the immune system, chloride, bromide, iodide, and the pseudo-halogen thiocyanate act as substrates for heme haloperoxidases-catalyzed reactions, producing reactive halogen species that contribute to host defense (*31*). Historically, the first characterized case of biological halogenation in humans was thyroglobulin iodination in the thyroid gland, leading to hormone synthesis (*32*). Since then, a few halogenated proteins have been identified in humans, for example, chlorinated apolipoprotein A1 in atherosclerotic plaques (*33*), brominated collagen IV (*34*), chlorinated plasma proteins (*35*) and histones (*36*). However, although these modifications have been detected, their functional or physiological roles in mammals systems have largely remained speculative. In this broader context, the identification and characterization of halogenated HNP1-3 represents, to our knowledge, the first demonstration of functionally halogenated proteins in human immunity. Our human diseases-derived haloproteome analysis points to a broader functional role for halogens in human biology. For instance, it raises the intriguing possibility that deiodinase enzymes (*37*), which regulate thyroid hormone metabolism, might also interact with iodinated HNP1-3, thus modulating iodine biological availability.

Through complementary biochemical, proteomic, and cellular analyses, we demonstrate that upon microbial or chemical activation, neutrophils generate both reactive oxygen and halogen species within intracellular compartments, leading to selective halogenation of HNP1–3 at conserved tyrosine residues. This modification is evolutionarily conserved in humans and rats, both of which express functional myeloid α-defensins, but is absent in mice, where the corresponding genes exist only as non-functional pseudogenes (*13*). Structural and proteomics analysis revealed preferential chlorination at Tyr21 and iodination at Tyr16 within hydrophobic pockets, a spatially precise modification tightly coupled to the oxidative burst. In clinical and pathological contexts characterized by neutrophil-driven inflammation, including cystic fibrosis, *S. aureus* abscesses, and *S. pneumoniae* severe pneumonia, halogenated HNP1–3 were abundantly detected, confirming their physiological relevance. In addition to HNP1–3, our study revealed numerous other proteins that undergo significant halogenation in pathological contexts. Especially noteworthy is the iodination of calprotectin, a critical antimicrobial protein that limits microbial growth by chelating iron and regulating *C. albicans* infections (*38*). Gene ontology further linked halogenated proteins to inflammatory signaling, damage repair and host defense pathways, highlighting their potential systemic immunological impact.

Despite the covalent addition of halogen atoms, HNP1s preserved their overall tertiary structure, maintaining the integrity of their hydrophobic cores while exhibiting increased surface hydrophobicity, a change that enhanced biological activity. Functionally, halogenated HNP1s displayed markedly improved toxin-neutralizing capacity, destabilizing bacterial virulence factors such as Anthrax Lethal Factor, Pneumolysin, and Leukocidins, thereby conferring robust cytoprotection to macrophages and erythrocytes. Beyond antitoxin activity, halogenation endowed defensins with amplified immunomodulatory properties. *In vitro*, halo-HNP1s strongly promoted neutrophil and monocyte chemotaxis through GPCR-dependent ERK1/2 signaling and secretion of the chemokines CCL2 and CCL3. Consistently, single-cell transcriptomic profiling of human PBMCs revealed that halo-defensins induced selective activation of monocytes, characterized by transcriptional upregulation of several chemokines (CCL2, CCL7, CXCL5), receptors (CCR1, IL1R1), interferon-related genes, and inflammatory mediators. Gene set enrichment analyses further highlighted activation of chemokine and immune signaling cascades, indicating that halo-HNP1s may bridge innate and adaptive immunity by promoting T-cells recruitment and priming. *In vivo*, these observations were recapitulated in a rat peritoneal inflammation model, where halo-RNP1s triggered robust infiltration of granulocytes and CD11b^high^ CD68^+^ monocytes, validating their immunostimulatory role.

Furthermore, our study demonstrates that neutrophils in pathological contexts generate hypoiodous acid (HOI), a reactive species long regarded as biochemically possible but physiologically irrelevant due to presumed scarcity of iodide in tissues (*5*). Given that HOI is over 100-fold more potent than HOCl in microbial killing (*39*), this finding may have implications for the understanding of host-pathogen interactions and micronutrient–immune system crosstalk. Indeed, epidemiological data showing that mild iodide deficiency impairs immune function, reversible upon supplementation, reinforce the physiological significance of this pathway (*40*,*41*,*42*).

In conclusion, our study establishes protein halogenation as a biologically meaningful post-translational modification that reshapes the function of antimicrobial proteins. By chemically integrating oxidative and non-oxidative antimicrobial pathways, neutrophils utilize HOX-mediated halogenation to reprogram HNP1-3 into multifunctional immune signaling molecules. This mechanism reveals an unexpected chemical layer of immune regulation and introduces halogenation as a previously unrecognized post-translational modification of innate immunity. This mechanism not only advances our understanding of neutrophil biology but also opens new avenues for designing peptide-based therapeutics that harness post-translational modifications for host defense.

## Acknowledgements

We thank Christian Frese from the Proteomics Research Platform at the Max Planck Unit for the Science of Pathogens for proteomic analysis. We thank the beamline staff at DESY/EMBL Hamburg, P14 for support with x-ray data collection.

This work was funded by the Max Planck Society (A.Z.). The structural analyses were funded by the NMR Based Structural Biology group, Max Planck Institute for Multidisciplinary Sciences (C.G.), Göttingen, Germany. Miha Milek was funded by the Deutsche Forschungsgemeinschaft (DFG) collaborative research center SFB 1588, Project number 493872418.

## Author Contributions

AF and AZ designed the study

AF, RH, KTT, MW, AB, UAA and PF performed experiments and analyzed the data SB solved the crystal structure of the chlorinated Cl21Y-HNP1 peptide, CG, VSM

DH, ST, AP and BS cared for the patients

AF, MR and AZ have interpreted the data and revised the manuscript for important intellectual content

AF wrote the manuscript

NMR analysis, figure preparation, and related writing were performed by VSM. SB solved the crystal structure of chlorinated Cl21Y-HNP1. CG discussed the NMR results and edited the manuscript.

## Data availability

The datasets generated and/or analyzed during the current study, including raw and processed data, are available from the corresponding authors upon reasonable request. Any additional materials, protocols, or code used in the study are also available upon request to ensure reproducibility of the results.

## MATERIALS AND METHODS

### Chemicals

All reagents were purchased from common vendors like Sigma Aldrich, Thermo Fischer Scientific or VWR Deutschland unless otherwise stated.

### Isolation of human neutrophils

The ethics council of the Charité Berlin (Germany) approved blood sampling and all donors gave informed consent according to the Declaration of Helsinki. Human neutrophils were isolated by a two-step density separation as described (*43*). Briefly, peripheral blood was layered on an equal volume of histopaque 1119 and centrifuged at 800 g for 20 min. PBMCs and neutrophil layers were collected separately, washed with PBS 0.2 % HSA and pelleted at 300xg 10 min. The pellet was resuspended in PBS 0.2 % HSA, layered on a discontinuous Percoll gradient (85%-65% in 2ml layers) and centrifuged at 800xg for 20 min. The neutrophil containing band was washed in PBS 0.2 % HSA and pelleted for 10 min at 300xg. The cell were counted with a CASY cell counter (OMNI Life Science).

### Isolation of human monocytes and differentiation into macrophages

Monocytes were isolated by positive selection (Miltenyi) according to the manufacturer’s instruction. Purity was determined by FACS staining with an anti-CD14 antibody. Monocytes were cultured in RPMI (Gibco, Carlsbad, USA) at 37°C and 7 % CO_2_. To generate monocyte-derived macrophages (MDM), monocytes were incubated in RPMI with phenol red supplemented with 5 ng/ml hM-CSF (Gibco) and hGM-CSF (Gibco) and incubated at 37°C and 7 % CO_2_ for 7 days. The medium was changed after three days.

### Cell lines

Adherent RAW 264.7 cells (ATCC, Manassas, USA) were cultured in RPMI-1640, 10% FCS (Hyclone®, Logan, USA), 1% penicillin-streptomycin and 5 mM glutamine. Cell cultures were maintained below 80% confluence. All cells were used at or below passage number 20 and incubated under standard conditions. Replicates are defined as cells harvested from different passages and/or flasks.

### Microbial culture

*S. aureus* strain USA300 was cultured in tryptic soy broth at 37°C for 18 h shaking. Bacteria were harvested, washed, and resuspended in PBS. *C. albicans* (clinical isolate SC 5314) and *C. albicans*^RFP+^ (*44*) were maintained on yeast peptone dextrose (YPD) agar plates or grown as previously described (*45*). *C. albicans* cells were washed twice in PBS, quantified by absorbance at 600 nm and confirmed by hemocytometer counting. *E. coli* XL1-Blue (Stratagene) was cultured overnight at 37°C in LB plus tetracycline, and subsequently subcultured to reach OD=0.4 (*46*).

### Isolation of mouse neutrophils

Mouse breeding and isolation of peritoneal neutrophils were approved by the Berlin state authority Landesamt für Gesundheit und Soziales under the approval number H0085/18. Animals were bred at the Max Planck Institute for Infection Biology in specific pathogen–free conditions, maintained on a 12-hour light/12-hour dark cycle, and food and water were provided ad libitum. C57BL/6 mice older than 12 weeks were injected i.p. with 1 ml of 7 % Casein solution in the evening and after 12 h. Three hours after the second injection, mice were sacrificed by cervical dislocation, the peritoneal cavity was flushed with 10 ml of PBS and the cells were collected. The cells washed three times with PBS, resuspended in 1 ml of PBS and neutrophils were separated by continuous density centrifugation for 20 min at 60000 g and the upper band containing the neutrophils was collected. Subsequently, the neutrophils were washed and counted in a CASY cell counter (OMNI Life Science). Purity was determined in cytospins stained with Giemsa; neutrophils were identified by their nuclear morphology.

### Isolation of rat neutrophils

Housing of rats were approved by the Berlin state authority Landesamt für Gesundheit und Soziales (ZH122). Rats were obtained from Envigo and kept at the Max Planck Institute for Infection Biology in Doppel Decker IVC on a 12-hour light/12-hour dark cycle and food/water ad libitum. 10-14 weeks old Sprague-Dawley rats were deeply anesthetized with isoflurane and blood was collected by final heart puncture. Neutrophils were purified by a modified mouse-neutrophil positive selection kit EasySep (Stemcell Technologies). The antibodies used for the negative selection were: mouse anti-rat CD32 (Becton Dickinson), mouse anti-rat-Ly76 (Miltenyi Biotech), APC conjugated mouse anti-rat antibodies: MHC class II, CD3, CD31, CD4, CD45RC, CD49d, CD8a (Miltenyi Biotech) and CD235a (LifeSpan Biosciences). After negative selection, the neutrophil-rich pellet was resuspended in erythrocytes lysis buffer and spun down for 10 min at 300xg. Purity was determined in cytospins stained with Giemsa; neutrophils were identified by their nuclear morphology.

### Patients

Blood samples were collected according to the Declaration of Helsinki with study participants providing written informed consent. CGD and MPO patients samples were collected with approval from the ethics committee Charité-Universitätsmedizin Berlin. The patients with X-linked chronic granulomatous disease (CGD) harbor mutations in the NADPH subunits genes (CYBB c.742dupA, CYBB c.868C > T, CYBB c.868C > T)^46, 47^. The ΔMPO donor bears a homozygous splice mutation (c.2031---2A >C/c. 20312 A>C, nomenclature according to (*47*,*48*), which generates null alleles which results in an immature protein (*48*).

Patients from sepsis and ARDS Registry (SPARE-14, Hannover Medical School) were treated on the intensive care unit and intubated for respiratory failure. Informed consent was given by the patient or a legal representative before intubation (#8146_BO_K_2018). BAL was performed preferably in the middle lobe or lingual or radiologically affected area. After sampling, macro-impurities were removed by filtration through sterile gauze. BAL samples were centrifuged at 500xg for 10min at 4°C and the supernatant (BAL fluid) was stored at -80°C until use.

Cystic Fibrosis patients were treated at the Charité-Universitätsmedizin Berlin and gave informed consent for the approved studies: BBL01a0041; RBB07a0039; RBB07a0373; RBB07a0371; RBB07a0414; RBB07a0469S. Sputum samples were collected after spontaneous expectoration or induced with inhaled hypertonic saline (sodium chloride 6%). Sample were cooled and processed within 24 hours.

Pus samples from *Staphylococcus aureus*-infected patients were obtained under the approval and guidelines of the Charité-Universitätsmedizin Berliń ethics committee (EA2/003/019). A 4-5 mm punch biopsy was taken from the abscess under local anesthesia. Informed consent was obtained from all subjects.

### Optical diffraction tomography and microscopy

Neutrophils (1×10⁵ cells per well) were seeded and stimulated with either phorbol 12-myristate 13-acetate (PMA, 100 nM) or *S. aureus* at a MOI=10:1 for 30 min. To visualize density distributions and fluorescence, we used a custom-built optical setup for correlative Optical Diffraction Tomography (ODT) and confocal fluorescence microscopy (*49*). Briefly, a 532 nm wavelength laser (MSL-III-532, CNI laser) illuminated samples seeded on imaging dishes (µ-Dish, ibidi). The complex optical fields were retrieved from spatially modulated holograms measured by illuminating samples from 150 different angles. By mapping the Fourier spectra of retrieved optical fields onto the surface of the Ewald sphere in the 3D Fourier space according to the Fourier diffraction theorem, 3D RI (Refractive Index) tomograms were reconstructed. Detailed principles for tomogram reconstruction can be found in previous reports (*50*). Custom-written MATLAB scripts (R2020a) allowed image acquisition, field retrieval and RI tomogram reconstruction. Along with ODT images, fluorescence image stacks were acquired using a Rescan Confocal Microscope(*51*). Neutrophil chromatin was stained with SPY-650 DNA (0.5 μM) and hypohalous acid with aminiphenyl fluorescein (APF, 5 μM). *S. aureus* cells were pre-labelled with DAPI (0.1 μM). For ODT we used a 60x water dipping objective (LUMPLFLN60XW, NA 1.0, Olympus Life Science) and a 100x oil immersion objective (UPlanFl, NA 1.3, Olympus Life Science). The 100x objective was also used for the confocal fluorescence imaging. We acquired fluorescence images with a Leica SP8 confocal microscope using an excitation wavelength of 488 nm, and the emission was filtered with a 505–550 nm filter.

### Neutrophil lysate preparation

Neutrophils (1-5×10^7^) were seeded in culture medium in microcentrifuge tubes at 1×10^7^ cells/ml. After spinning down for 10 min at 300xg, cells were resuspended in RIPA buffer with 20 µM neutrophil elastase inhibitor GW311616A (Biomol) and Halt protease inhibitor cocktail (PIC), mixed and incubated at 37°C with gentle rotation, before centrifugation at 1000xg (30 s) to collect the supernatant. We added freshly boiled 5 X sample loading buffer (50 mM Tris-HCl pH 6.8, 2% [w/v] SDS, 10% glycerol, 0.1% [w/v] bromophenol blue, 100 mM DTT), vortexed and boiled for 10 min with agitation before flash freezing in liquid nitrogen for storage at –80 °C. We used DPI and 4-ABAH inhibitors at concentration of 100 nM and 300 µM, respectively, and preincubated the cells for 30 minutes at 37°C.

### Halo-tyrosine analysis

Cell Lysates (1-5×10^7^ cells, after 1 h stimulation or naive) and amino acid standards (tyrosine and halo-tyrosines) were acid-digested in 1 ml vacuum-hydrolysis tubes in a total volume of 100 µl of 4M methanesulfonic acid and 0.1% phenol for 18 hours at 110 °C. The mixtures were neutralized with 4M NaOH to a pH of 4 and brought to a volume of 2 ml with 0.1% acetic acid. After centrifugation (10.000 x g 2 min), the supernatants were passed over polymeric strong cation exchange SPE-columns (Strata-X-C 33µm, 30 mg tubes, Phenomenex) and equilibrated with 0.1% acetic acid. After washing with 2 CV (column volume) of 0.1 % acetic acid, compounds were eluted with a 4% ammonia solution in 50% methanol. Eluates were dried in a SpeedVac evaporator and solubilized in 20µl of water.

We used the manufacturer’s protocol (Waters) for AccQ-Tag derivatization with some modifications. In brief, 6-aminoquinolyl-N-hydroxysuccinimidyl carbamate (AQC) was reconstituted in acetonitrile at 3mg/ml and heated at 55 °C for 2 min. 20µl of the AQC reagent were mixed immediately with 20µl of the solid-phase-extracted sample and 60µl 0.5M sodium borate pH 8.5. The mixture was incubated for 1min at room temperature, vortexed at 55°C for 15 minutes in the dark. 1 to 5µl were applied to UPLC-chromatography. To verify the presence of unbound chloro-tyrosine, cell lysates were analyzed directly, without undergoing acid hydrolysis. Derivatized amino acids were separated by reversed phase chromatography on an AccQ-TAG Ultra C18-UPLC column (2.1 x 100mm, 1.7µm, Waters) with a linear gradient from 0.1% formic acid in 10% acetonitril to 90% acetonitrile over 6 min at 55°C and a flow rate of 0.5 ml/min. Eluted compounds were detected by fluorescence (excitation and emission at 250 and 395 nm, respectively) and by ESI-MS detection (QDa, Waters). The QDa was operated in an electrospray positive ion mode by applying a voltage of 0.8 kV. The cone voltage was set at 15 V. The probe temperature was set at 600 °C. We acquired a full mass spectrum between m/z 100 and 1200 with a sampling rate of 2.0 points/sec.

### Ammonium sulphate fractionation

Human neutrophils (1-5×10^7^) were lysed (RIPA buffer) and centrifuged for 15 minutes at 2500xg, at room temperature and the supernatant further centrifuged at 5000xg for 15 minutes at room temperature. Proteins were sequentially precipitated from 1 mL of this lysate by stepwise addition of solid ammonium sulphate with stirring till saturation, followed by incubation on ice for 1 h and centrifugation at 10,000xg at 4°C for 15 minutes. After each centrifugation, the pellet obtained was resuspended in 1 mL of buffer containing 10 mM phosphate, pH 7 and repeated with 50, 60, 70 and 80% of ammonium sulphate saturation. Aliquots of precipitated fractions were analyzed in 4-12% sodium dodecyl sulfate polyacrylamide gel electrophoresis (SDS-PAGE).

### Liquid Chromatography−Mass Spectrometry Analysis (LC−MS)

For identification and relative quantification of proteins, gel pieces were tryptic digested, as described (*52*). Tryptic peptides were analyzed by LC-MS using a Q Exactive Plus mass spectrometer. Peptide mixtures were fractionated by an Ultimate 3000 RSLCnano with a two-linear-column system. Digests were concentrated for 4 min onto a trapping guard column (PepMap C18, 5 mm x 300 μm x 5 μm, 100Ǻ). Then, samples were eluted after 55 min from an analytical column (75 µm i.d. × 250mm nano Acclaim PepMap C18, 2 μm; 100 Å LC column). Samples were separated using a mobile phase from 0.1% formic acid (FA, Buffer A) and 80% acetonitrile with 0.1% FA (Buffer B). We used a 15 min active linear gradient from 3 to 53 % of buffer B at a flow rate of 250 nL/min. For 50 min the Q Exactive instrument was operated in a data dependent mode to automatically switch between full scan MS and MS/MS acquisition. Survey full scan MS spectra (*m*/*z* 350–1600) were acquired in the Orbitrap with 70 000 resolution (*m*/*z* 200) after 50 ms accumulation of ions to a 3e6 target value. Dynamic exclusion was set to 10s. The 10 most intense multiply charged ions (*z* ≥ 2) were sequentially isolated and fragmented by higher-energy collisional dissociation (HCD) with a maximal injection time of 200 ms, AGC 1e6 and resolution 17 500. Typical mass spectrometric conditions were as follows: spray voltage, 2.0 kV; no sheath and auxiliary gas flow; heated capillary temperature, 275°C; normalized HCD collision energy 27%.

Additionally, the background ions m/z 391.2843 and 445.1200 acted as lock mass. Identification and relative label-free quantification of the proteins and peptides were performed with MaxQuant software version 1.6.0.1 (*15*) using the following search parameter set: Spectra were matched to a human (20,386 reviewed entries, downloaded from uniprot.org) or a rat (8,071 reviewed entries, downloaded from uniprot.org), a contaminant, and decoy database, enzyme: trypsin/P with two missed cleavage, static modification: carbamidomethylation (C), variable modifications: protein N-acetylation oxidation (M), dichlorination and chlorination, bromination, diiodination, triiodination and iodination (Y), mass tolerances for MS and MSMS: 10 ppm and 0.02 Da, and a calculated peptide FDR 1%.

### 36Cl#radiolabeling

Customed RMPI 1640 medium was enriched with NaCl containing ^36^Cl radioactive isotope (Chlorine-36 radionuclide as potassium chloride / s.A. 2-20 mCi/g Cl, Conc. 0.1 mCi/ml, Aqueous solution, M.W. 74.5, Biotrend). Freshly isolated neutrophils (1-5×10^7^) were incubated for 1 h in the ^36^Cl enriched medium (25 uCi) before adding PMA (100 nM) or *S. aureus* (MOI 10:1). After 1 h of stimulation, cells were harvested and lysed in RIPA buffer. Whole lysate was resolved by SDS-PAGE and then the gel was dried and exposed to a radio-sensitive film for 10 days.

### Total ROS assay

To assess total ROS production, 10^5^ neutrophils were incubated in bicarbonate-free Seahorse medium (Seahorse XF Media, Agilent) for 25 minutes in a CO_2_ free incubator. After it, the cells were pretreated with luminol (50μM), and activated with 50nM PMA. ROS production was measured by monitoring luminol luminescence for 3 hours as previously described (*3*).

### Hypohalous acids measurement

Human neutrophils or monocytes, as well as mouse and rat neutrophils were seeded (10^5^ cells per well) onto a glass-bottomed dish and incubated with APF (5 μM) for 30 min at 37°C before stimulation with PMA (100 nM), *S. aureus* (MOI 10:1), or hydrogen peroxide (1 mM). The fluorescence was monitored using a Fluoroskan Ascent fluorescence spectrometer at an excitation wavelength of 488 nm and an emission wavelength of 550 nm for 3 hours. We acquired fluorescence images 10 min before, and 10, 30 and 60 minutes after stimulation with PMA and *S. aureus* with a Leica SP8 confocal microscope using an excitation wavelength of 488 nm, and the emission was filtered with a 505–550 nm filter.

### Phagocytosis analysis

We determined phagocytosis activity as previously described (*53*); freshly isolated neutrophils (10_6_) were incubated with opsonized bacteria in RPMI with 0.1% human serum albumin in a bacteria to neutrophils ratio of 10:1.

To quantify phagocytosis rate of macrophages, opsonized (human serum) *C. albicans*^RFP+^ was coincubated with HMDM (10^6^) at MOI 1:1 for 4h in RPMI medium at 37°C. After it, samples were analyzed microscopically and images processed by ImageJ. Phagocytic rate was calculated by automatic segmentation using the software Fiji.

### UPLC and hydrophobicity analysis

Peptides (10-50 µg) were dissolved in 0.1% trifluoroacetic acid (TFA) and injected onto a BEH C18 reverse phase column (2.1 mm x 100mm, 1.78µm, Waters) using a linear gradient from 10% to 80% acetonitrile (containing 0.1% TFA) within 6 min at a flow rate of 0.5 ml at 45°C. Peptides were detected at 280nm. The nile red hydrophobicity assay was previously described (*54*) was dissolved in DMSO to 0.25 mM. We mixed 10-50 µg of HNP1, ClY21-HNP1 and IY16-HNP1 in PBS, pH 7.4, with 0.25 μM Nile red and monitored fluorescence with a Fluoroskan Ascent fluorescence spectrometer (Agilent) at 550 nm excitation and 570–700 nm emission. Data were fitted with GraphPad/ Prism.

### HNP1 peptides synthesis

Chemical synthesis of HNP1 and halogenated analogs was performed by Biosynth International, Inc. (USA, MA). Machine-assisted solid phase chemical synthesis of HNP1 and its analogs, was performed by Biosynth using the 2-(1H-benzotriazolyl)-1,1,3,3-tetramethyluronium hexafluorophosphate activation/N,N-diisopropylethylamine *in situ* neutralization (*55*).

### X-ray crystallography

We dissolved synthetic Cl21Y-HNP1 peptide (20 mg/ml) concentration in distilled water and obtained crystals by the vapor-diffusion technique with sitting drops. 100 nl of protein solution were mixed with 100 nl of well solution using a grid screen as published (*56*) (0.1 M imidazole, 1.0 M sodium acetate trihydrate, pH 6.5). Crystals grew within one week and were cryoprotected by transferring them for one minute to well solution supplemented with 25 % glycerol and flash-cooled by plunging them into liquid nitrogen. Diffraction data were collected at the EMBL beamline P14 at PETRA III, DESY, Hamburg, Germany. Data were processed and analyzed with autoProc (*57*) and data collection statistics are shown in Table S1. The structure was solved by molecular replacement with PHASER (*58*) using the crystal structure of wildtype HNP1 (PDB accession code: 3gny) as search model. Refinement was performed initially with Refmac (*59*) alternating with manual model building in Coot (*60*). For a final refinement we used phenix.refine (*61*). Refinement statistics are shown in Table S2. The coordinates of the Cl21Y-HNP1 structure were deposited to the protein data bank (PDB accession code: 9fqb).

### Structural alignments

Superposition of the structural models was performed by secondary-structure matching with SSM (*62*) of protein backbone C^α^ atoms as implemented in Coot (*60*).

### NMR Spectroscopy

Protein samples of ClY21-HNP1 and IY16-HNP1 were transferred in 5 mm NMR tubes and analyzed at 333.2 K using a Bruker 900 MHz Avance NEO spectrometer equipped with a cryogenically cooled probe. Two-dimensional (2D) ^1^H–^1^H TOCSY and NOESY spectra were acquired with 4096 points in the direct dimension (F2; spectral width: 14.20 ppm; acquisition time: 180.22 ms) and 512 points in the indirect dimension (F1; spectral width: 14.20 ppm; acquisition time: 22.52 ms). The NOESY spectra were collected with a mixing time of 500 ms. In addition, (2D) ^1^H–^13^C HSQC spectra were acquired using 4096 points in F2 (spectral width: 13.01 ppm; acquisition time: 196.61 ms) and 256 points in F1 (spectral width: 165.00 ppm; acquisition time: 3.85 ms). All 2D datasets were processed with zero-filling to 8192 and 2048 points in the F2 and F1 dimensions, respectively, and apodized using a sine square window function with a sine bell shift (SSB) of 2 in both dimensions. All one-dimensional (1D) and two-dimensional (2D) NMR spectra were processed using TopSpin 4.0.8 (Bruker Biospin) and analyzed with CCPNMR Analysis V3 (http://www.ccpn.ac.uk/ccpn).

### HNP1 killing assay

HNP1, ClY21-HNP1 and IY16-HNP1 at concentrations between 1 to 100 μg/ml were diluted in 10 mM sodium phosphate (Na_2_HPO_4_) solution at pH 7.4, and incubated with bacteria (1-5×10^5^). We included vehicle as well as HNP1s denatured with 10 μM DTT as controls. After 1 h at 37°C we sampled at the indicated time points and plated.

In *C. albicans* experiments, HNP1, ClY21-HNP1 and IY16-HNP1 were diluted in 10 mM sodium phosphate (Na_2_HPO_4_) solution at pH 7.4, with concentrations ranging from 1 to 100 μg/ml. After incubation with *C. albicans* (10^4^) for 2 h at 37°C, a sample was plated on YPD plates. Additionally, we measured *C. albicans* killing by HNP1 with the by XTT metabolic assay as previously reported (*63*).

### Chemotaxis

Human neutrophils (1×10^5^) and monocytes (1×10^5^) were seeded in the upper changed of multiwell chambers (Cell Migration/Chemotaxis Assay Kit, Abcam) with a 5 (for monocytes) or 3 (for neutrophils) µm pore size filter while the lower chamber contained chemotactic factors. After 1 h incubation at 37°C, the filters were removed, and the migrated cells stained with DRAQ5. Using a hemocytometer microgrid (*64*), we counted the number of cells that migrated to the lower chamber toward the chemoattractant. fMLP at 50 nM (for neutrophils) or CCL2 at 50 ng/ml (for monocytes) were used as positive controls and the results expressed as total number of cells migrated through the pores.

### Thermostability assay

For each condition, 10 μL of protein sample were prepared in PBS at 0.4 mg/ml. The protein samples were loaded into UV capillaries on Tycho NT.6 (NanoTemper Technologies) for label-free thermal shift analysis. Unfolding profiles we measured the change of fluorescence emission ratio (350/330nm) over a temperature range from 35 to 95°C.

### Anti-toxin experiments

Human neutrophils, primary human monocytes or RAW 264.7 cells were seeded in a 96-well plate at a density of 1×10^5^ cells per well in RPMI with 5% human serum. We added 20 nM LF, 50 nM PA, 50 nM LLO, 50 nM PLY or 50 nM of the PVL components LukS and LukF in equimolar ratio, and 0.1 to 10 μM of HNPs simultaneously to the cells. We determined cell viability by LDH release 1 and 2 hours later.

### Cytotoxicity assay

Human neutrophils, monocytes, HMDM and RAW 264.7 cells were seeded in RPMI in 6 well plates with a density of 1×10^6^ cells/ml. Inhibitors were preincubated with cells for 30 min at 37°C at the indicated concentrations. We used CytoTox 96 Non-Radioactive Cytotoxicity Assay (Promega) to measure toxicity after 1 or 2 hours incubation.

### LF metalloproteolytic activity

We used an N-acetylated, C-7-amido-4-methylcoumarin (AMC) derivative of a 14-mer fluorogenic peptide substrate (Merck) designed from the MEK-2 template to measure Anthrax lethal factor (LF) proteolytic activity. We tested inhibition with HNP1, ClY21-HNP1 and IY16-HNP1 at concentration between 0.1 to 10 µM. Proteolytic activity of LF was monitored using a Fluoroskan Ascent fluorescence spectrometer (Agilent).

### ELISA analysis of chemokines

Supernatant of HMDM (10^6^ cells per well) stimulated with HNP1, ClY21-HNP1, IY16-HNP1 (4 hours) were harvested and analyzed by ELISA (Human ELISA kit, Abcam) for CCL2 and CCL3, according to the instructions of the manufacturers. The supernatants of stimulated cells were used undiluted. For blocking of plates and dilution of standard curves, ELISA dilution buffer was used, unless it was specified differently in the instructions. ELISA plate were washed with PBS-T.

### Western blot

Cell lysate (human monocytes, 10^6^) samples for western blotting were reduced in 1× LDS sample buffer (Invitrogen) and DTT was added at a final concentration of 100 mM before boiling at 70°C for 15 min. The samples were run at 120 V for 1.5 h in MES running buffer (Invitrogen) using the NuPage Invitrogen Mini gel tank system in precast 4–12% gradient Bis-Tris gels. The gels were then directly stained with Instablue (Abcam) or transferred using a BioRad wet tank system onto a 0.22 µm PVDF membrane (Amersham). After transfer, the membranes were blocked for 1 h in 5% non fat-dried milk followed by primary antibody overnight incubation at 4°C (Cell Signaling Technology, p44/42 MAPK (Erk1/2) Antibody #9102 1:1,000; Cell Signaling Technology, Phospho-p44/42 MAPK (Erk1/2) (Thr202/Tyr204) 1:1,000) and then a secondary HRP conjugated antibody (Jackson Labs 1:20,000) for 1 h before washing, and the bands were developed with ECL (Pierce) using the Bio-Rad ChemiDoc. As control, human monocytes were pre-incubated with *Bordetella pertussis* toxin (Sigma) at concentration of 100 ng/ml for 1 hour.

### Hemolysis

Red blood cells (RBCs; human, RBC9, Sigma) were washed four times with PBS and resuspended in PBS at 5% (v/v). LLO or PLY and HNP1 peptides were preincubated on ice for 15 min before adding RBCs to 96-well plates, with a final volume of 0.1 ml per well. After gentle mixing, hemolysis was monitored at room temperature using a VersaMax microplate reader. Absorbance at 700 nm (A_700_) was recorded every 30 s for 30 min, with automatic plate shaking before each reading. Hemolysis protection by HNP1s was quantified by plotting the change in A_700_ (ΔA_700_).

### Opsonophagocytic Killing Assay (OPK)

Neutrophils from healthy donors were isolated from buffy coats (Gulf Coast Regional Blood Center) using a Ficoll gradient and suspended in RPMI. We pre-coated 96-well tissue culture with 20% pooled human serum (SeraCare #1830-0003) for 30 min at 37°C, washed twice, and primed neutrophils with 10 µL GM-CSF (36 µM; Gibco #PHC2015), 10 µL HPN1s (1 µM final in PBS), or mock PBS for 30 min at 37°C 5% CO_2_. We infected these neutrophils with subculured *S. aureus* subcultures mixed with 50 µL of pooled human serum at a multiplicity of infection of 5-1 (bacteria to neutrophils). After a 3.5 h incubation we lysed the neutrophils with 1% saponin and added to the filter plates (USA Scientific #2920-1000) containing 100 µL TSB 0.1% TCC, centrifuged at 100 x g for 1 min and incubated at 37°C overnight in a sealed bag. Colonies were fixed in 70% ethanol, dried, and imaged on a CTL ImmunoSpot analyzer and quantified with ImmunoSpot SC Suite software (v7.0).

### scRNAseq data processing

Raw sequencing reads was processed with Cellranger multi v8.0.0 (10X Genomics) and included alignment to the human genome (GRCh38), probe barcode demultiplexing and generation of count matrices with mRNA read counts. Ambient RNA background was removed using CellBender remove-background (v0.3.0) starting from demultiplexed raw count matrices. The filtered count data was then processed with Seurat (v5.1.0) (*65*, *66*). We retained cells that had lower than 10% mitochondrial genes, had more than 500 and less than 6000 quantified genes. We recovered 200741 cells with a median of 2244 quantified genes per cell.

During exploratory quality control, donor 3 (vehicle and HNP1 conditions) was identified as an outlier. Pseudo-bulk gene expression profiles from this donor showed opposite fold-change directionality compared with the other donors, and PCA and sample–sample correlation analyses consistently separated donor 3 from the remaining samples. Normalization and batch-correction approaches (Seurat integration) did not resolve the discrepancy, indicating a donor-specific technical anomaly. Therefore, donor 3 was excluded from downstream analyses.

### Cell annotation

We annotated the cell types with Seurat multimodal reference mapping workflow using the human PBMC atlas. Briefly, we normalized and scaled the counts using SCTransform, followed by FindTransferAnchors and MapQuery functions. Three layers of granulation in the reference were transferred to the query data to assign each cell to a reference type. We embedded cell into the reference space using IntegrateEmbeddings. We obtained scores for the predicted cell type for each granularity level, which served as quality control of the cell assignment. We only used granularity levels 1 and 2 for downstream analysis. Level 3 was excluded because of its low granularity scores.

### Differential expression and gene set enrichment analyses

We summed the gene-wise read count for each sample over all cells and for each cell type at granularity level 1 and then analyzed differential pseudobulk expression with DESeq2 (*67*) using the model design ∼batch + group. We extracted pairwise comparisons between different groups from the same results object, to consider the total variance from all samples in the experiment. We used tmod (*68*) to perform gene set enrichment analysis on the DESeq2 results. The gene list sorted by DESeq2 P value was the input used into the tmodCERNO test function. As a threshold for tmod results, which outputs adjusted P value, as well as effect size (AUC), we considered enriched pathways to have an AUC>=0.70, and an adjusted P value < 0.0001, unless specified otherwise in the figure legends. To define genes with increased or decreased expression we used a cutoff of adjusted P value <0.05 and absolute log2-transformed fold change of >0. The immune related gene sets for pathway enrichment, were provided by the tmod package (*68*,*69*,*70*)

### *In vivo* peritoneal inflammation model in rats

Adult female Sprague Dawley rats, bred and tested from Selvita LTD (Zagreb, Croatia) were randomly allocated to experimental groups and acclimatized for at least 1 week. Rats were injected intraperitoneally with 10 μg of peptides or vehicle (water) in a total volume of 1 ml per animal. After 24 h, the rats were sacrificed and the peritoneum lavaged with 50 ml of cold PBS. The cells were washed and fixed for flow cytometry.

## Statistical analysis

Statistical analyses were done using GraphPad Prism v10 software and the R statistical environment. All statistical details of experiments can be found in the figure legends.

## SUPPLEMENTARY FIGURES

**Figure S1.**
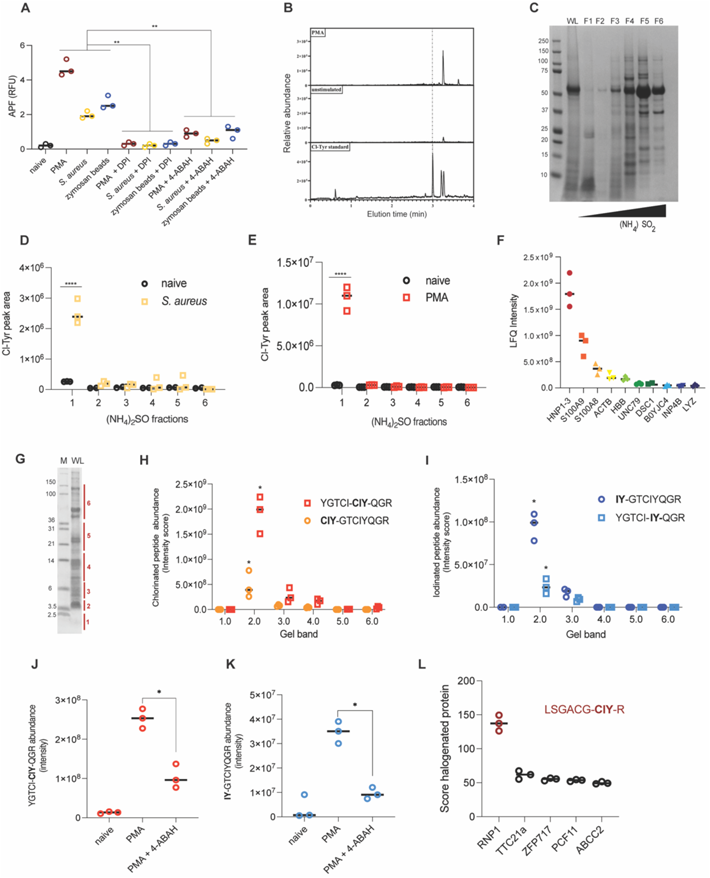
Neutrophils Halogenate α-Defensin 1–3. (A) Human neutrophils treated with diphenyliodonium (DPI) or 4-aminobenzoic acid hydrazide (4-ABAH, 300 µM), before stimulation with Phorbol-12-myristate-13-acetate (PMA, 100 nM), S. aureus (MOI=10:1) or zymosan (10 µg/mL) as indicated. Hypohalous acids was measured with 3′-(p-aminophenyl) fluorescein (APF) 60 minutes after stimulation. **p < 0.001, two-way ANOVA with Tukey’s multiple comparisons test. N=3. (B) Representative ESI-MS elution profile of non-hydrolized lysates from naive and PMA stimulated human neutrophils. Chloro-tyrosine standard was used as control (dashed line); N=2. (C) SDS-PAGE of the ammonium sulphate precipitation fractions of PMA stimulated human neutrophil stained with Coomassie blue. WL = whole lysate. (D-E) Representative ClY quantification of the ammonium sulphate fractions of human neutrophils activated with S. aureus (D) or PMA (E). Data points indicate technical replicates. Data were analyzed by one-way ANOVA with Bonferroni multiple comparisons test; ****p < 0.0001. N=4. (F) LC-MS quantification of the proteins in fraction 1 from panel C in this figure. N=3. (G) Representative tricine 16% denaturating acrylammide gel of PMA-stimulated human neutrophils cell lysate. N=3. The bands analysed in panels H and I are indicated and labeled in red. (H-I) Representative LC-MS analysis of chlorinated (H) or iodinated (I) HNP1-derived peptides extracted from the tricine 16% gel bands in panel G. Data points indicate technical replicates. Data were analyzed by one-way ANOVA with Bonferroni multiple comparisons test; *p< 0.01. N=3. (J-K) Representative mass spectrometry-based quantification of chlorinated (J) and iodinated (K) HNP1-3 derived peptides from human neutrophils (2×10⁵ cells) in the presence or absence of 4-ABAH before stimulation with PMA. Data points indicate technical replicates. Data were analyzed by one-way ANOVA with Bonferroni multiple comparisons test; **p< 0.01; ***p< 0.001. N=3. (L) Representative LC-MS analysis of chlorinated peptides from S. aureus stimulated rat neutrophils. Amino acid sequence indicated that tyrosine 22 is chlorinated. N=3.

**Figure S2.**
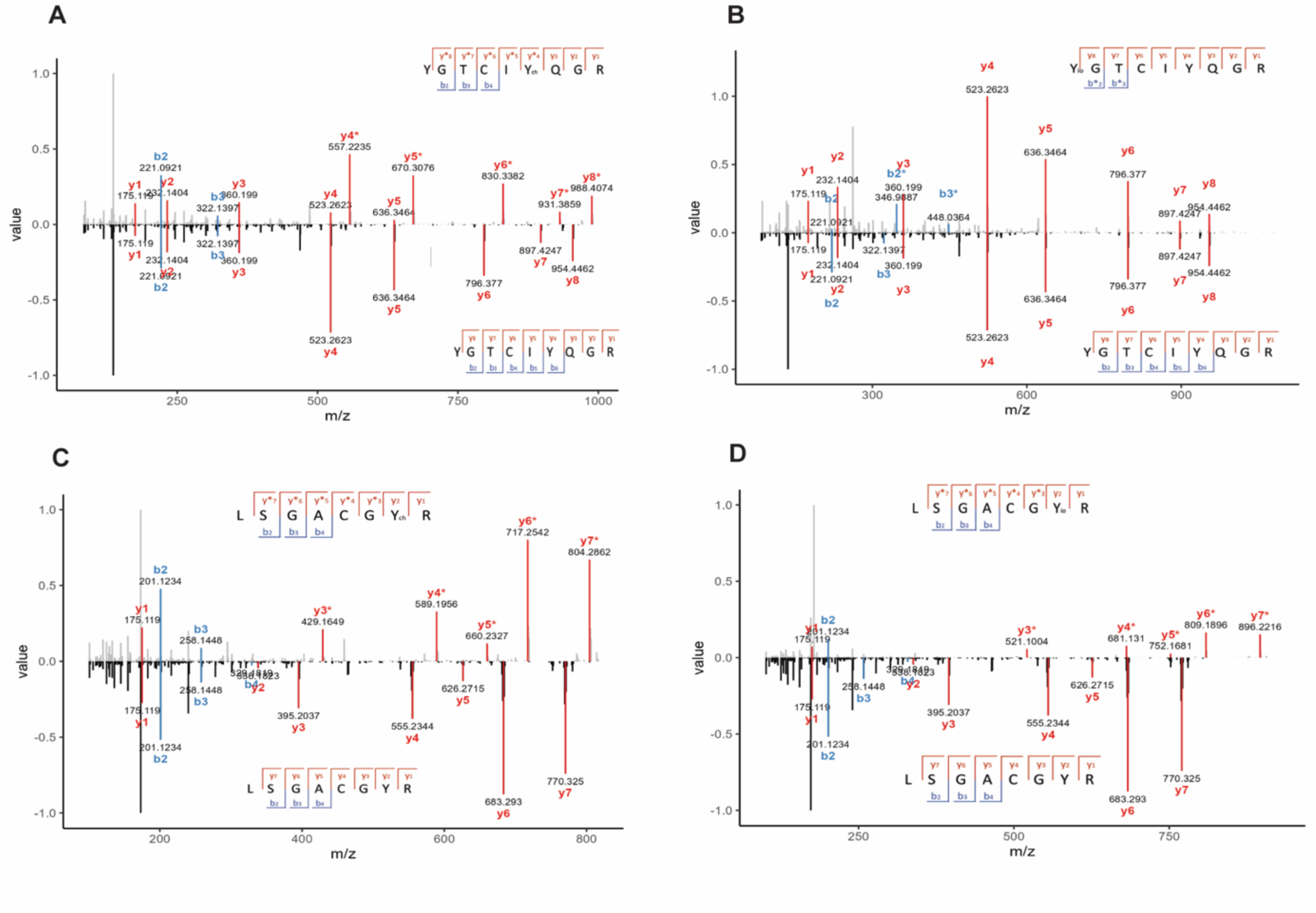
MS/MS spectra of halogenated HNP1-3 and RNP1-2 peptides. Chlorination (+33.9610 Da) or iodination (+125.8966 Da) of tyrosine residues causes a mass shift in the corresponding b- or y-ions compared to the unmodified peptides. (A) Chlorination of HNP1-3 peptide YGTCIYchQGR: the y-ions 4-8 (red) in the top spectrum show the m/z values for the peaks for y4-y8 indicate the mass shift of +33.9610 Da (highlighted as *) compared to the corresponding fragment ions in the bottom spectrum. (B) Iodination of HNP1-3 peptide YioGTCIYQGR: the b-ions 2-3 (blue) in the top spectrum show the m/z values for the peaks for b2-b3 indicate the mass shift of +125.8966 Da (highlighted as *) compared to the corresponding fragment ions in the bottom spectrum. (C) Chlorination of RNP1-2 peptide LSGACGYchR: the y-ions 3-7 (red) in the top spectrum show the m/z values for the peaks for y3-y7 indicate the mass shift of +33.9610 Da (highlighted as *) compared to the corresponding fragment ions in the bottom spectrum. (D) Iodination of RNP1-2 peptide LSGACGYioR: the y-ions 3-7 (red) in the top spectrum show the m/z values for the peaks for y3-y7 indicate the mass shift of +125.8966 Da (highlighted as *) compared to the corresponding fragment ions in the bottom spectrum.

**Figure S3.**
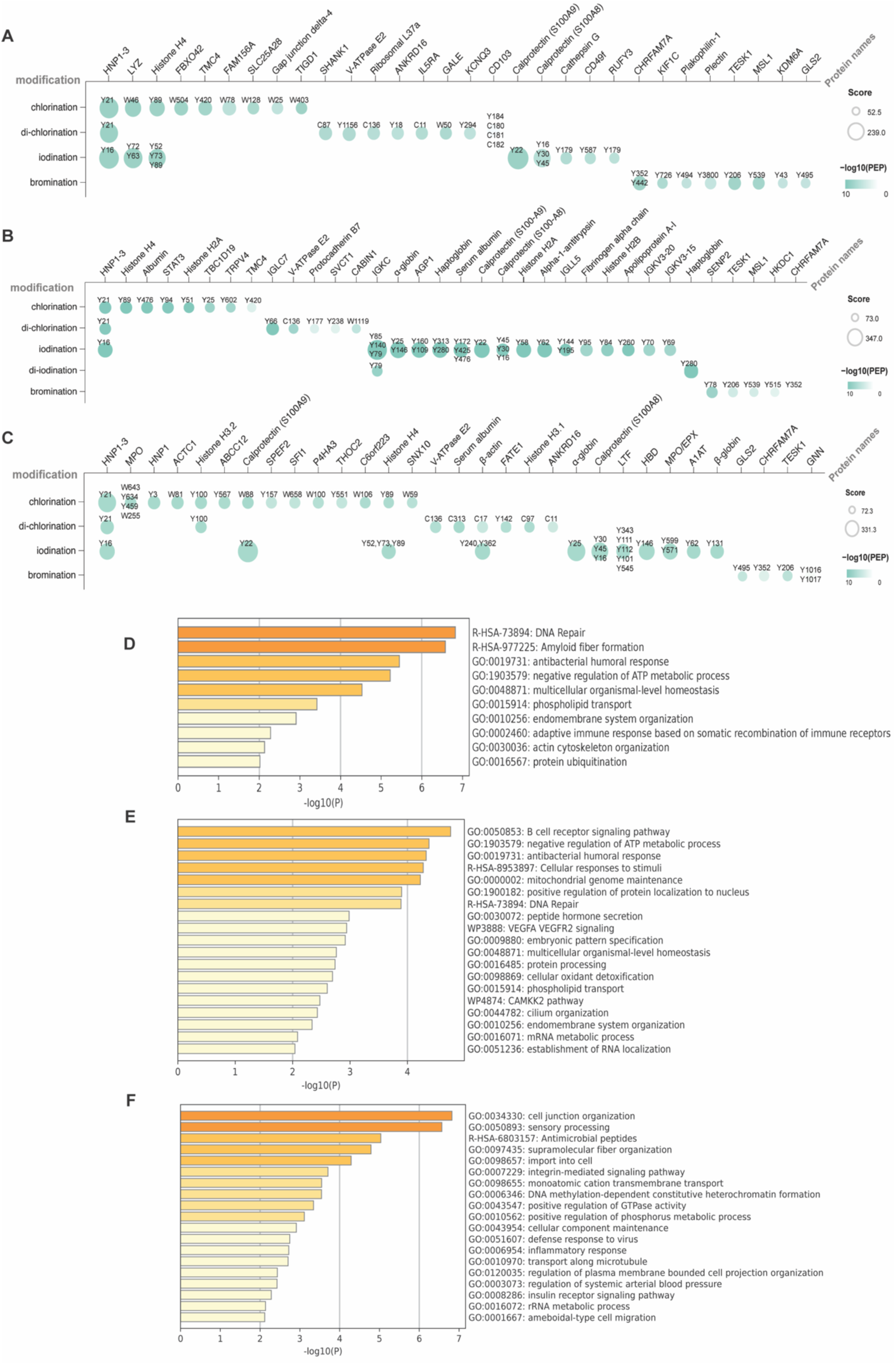
The haloproteome of patients reveals HNP1-3 as major halogenation target. (A-C) Mass spectrometry analysis of the haloproteome of (A) cystic fibrosis (CF) patients sputum samples, (B) S. pneumoniae infected patients BAL and (C) S. aureus skin abscess. The analysis displays proteins with andromeda score >50, type of halogenation and aminoacid residues. Score indicates andromeda score and PEP indicates posterior error probability. (D-F) Pathway enrichment analysis of halogenated proteins identified in CF sputum (D), S. pneumoniae BAL (E) or S. aureus abscess (F) samples. Enriched functional categories were identified using differential overrepresentation (DO) clustering. The identifiers displayed in the plots originate from distinct biological databases that are commonly integrated in enrichment pipelines: GO:entries correspond to Gene Ontology (GO) terms, which capture broad biological processes, molecular functions, or cellular components; R-HSA-identifiers denote Reactome pathways, which represent curated and mechanistic pathway maps specific to Homo sapiens; and WPentries correspond to WikiPathways, a community-curated collection of pathway maps.

**Figure S4.**
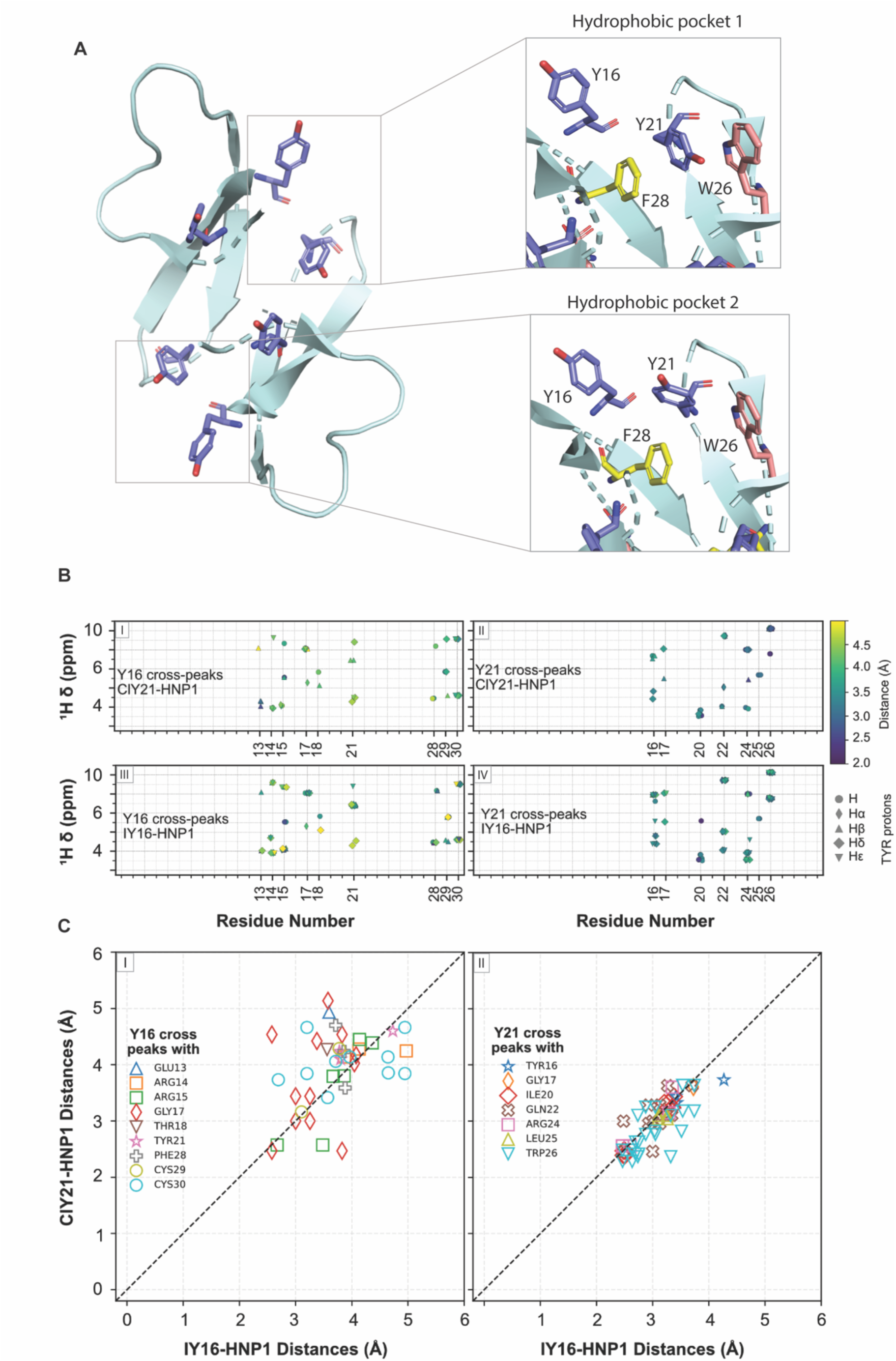
Halogenation fine-tunes HNP1-3 physicochemistry without structural disruption. (A) 3D-structure of HNP1 highlighting tyrosines (purple) and spatial organization of the hydrophobic pocket 1 and 2, including phenylalanine 28 (yellow) and tryptophan 26 (red), with the corresponding amino acids (PDB:1gny) generated with PyMOL Molecular Graphics System. (B) NOESY cross-peaks between the protons of Tyr16 (I and III) and Tyr21 (II and IV) with protons from other amino acids in the HNP1 structure. Data from ClY21 and I16Y are shown in panels (I and II) and (III and IV), respectively. The X-axis denotes the residue number of the amino acids exhibiting cross-peaks with the specified tyrosine residue, while the Y-axis represents the chemical shift of the corresponding partner proton. (C) Cross-peak intensities for Tyr16 and Tyr21 were normalized relative to the G17(H)-Y16(Hα) (I) and Q22(H)-Y16(Hα) (II) interactions, respectively. These normalized values were subsequently converted to interatomic distances using the relation:

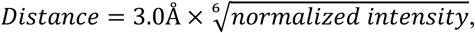

and are displayed using the key with different colors and symbols.

**Figure S5.**
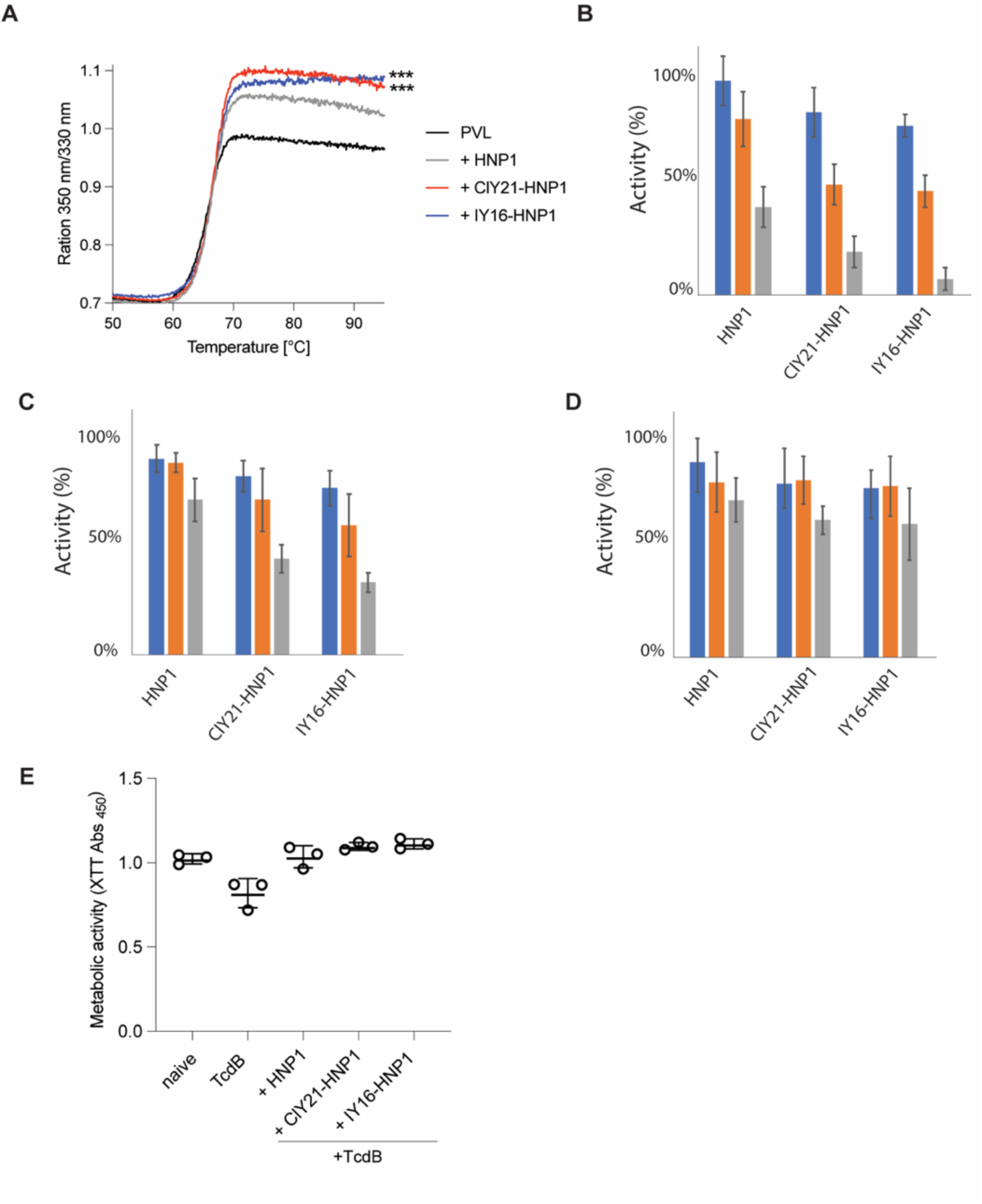
Anti-toxin activity of halogenated HNP1s. (A) Thermostability assay assessing the binding between Panton–Valentine Leukocidin (PVL) to HNP1, ClY21-HNP1 or IY16-HNP1. Data were analyzed by two-way ANOVA with Tukey’s multiple comparisons test. N=3, ***p < 0.001. (B-D) Anti-toxin activity measured as inhibition of neutrophil cell death (LDH release) of HNP1, ClY21-HNP1 or IY16-HNP1 against the *S. aureus* pore-forming toxins HlgAB (B), HlgCB (C), and LukAB CCR45 (D). The concentrations tested were 0.1 µM (blue), 1 µM (orange), and 10 µM (gray). Bars represent means ± SEM from three independent experiments (N=3). (E) Representative anti-toxin activity measured as metabolic activity of human monocytes treated with 1 µM *Clostridium difficile* toxin (TdcB) in presence of HNP1, ClY2-HNP1-or IY16-HNP1 (N=3).

**Figure S6.**
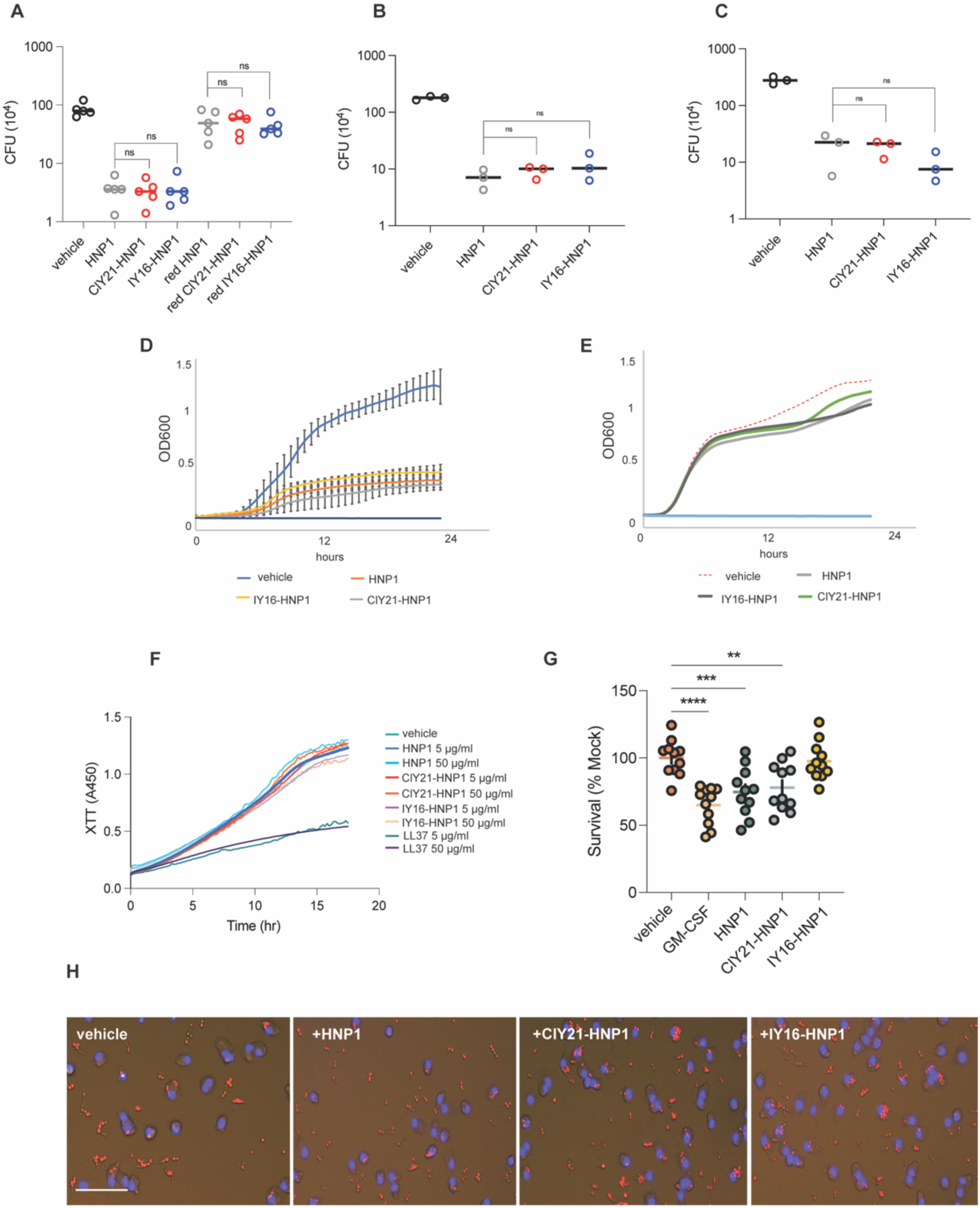
Microbial killing by halogenated HNP1s. (A) Colony-forming unit (CFU) assays to measure the antimicrobial activity of HNP1, ClY21-HNP1or IY16-HNP1 (50 µg/ml) against *S. aureus*. Peptides reduced with DTT (red HNP1, red ClY21-HNP1or red IY16-HNP1) were used as negative controls. Data represent means ± SEM (N=5). Statistical significance was assessed by one-way ANOVA with Dunnett’s multiple comparisons test; ns indicates non-significant differences. (B,C) Colony-forming unit (CFU) assays to measure antimicrobial activity as described in A with *Escherichia coli* (B), or *Candida albicans* (C) at a concentration of 50 µg/ml, N=3. Statistical significance was assessed by one-way ANOVA with Dunnett’s multiple comparisons test; ns indicates non-significant differences. (D, E) Growth curves of *S. aureus* cultured in RPMI (D) or TSB (E) in the presence of HNP1, ClY21-HNP1or IY16-HNP1 (50 µg/ml). Data are shown as means ± SEM from three independent replicates (N=3). No significant differences were observed. (F) Metabolic activity of *C. albicans* cultured in TSB in the presence of HNP1, ClY21-HNP1or IY16-HNP1 (5 and 50 µg/ml). HNP1 5 and 50 µg/ml (dark blue, light blue), ClY21-HNP1 5 and 50 µg/ml (red, orange) and IY16-HNP1 5 and 50 µg/ml (purple, pink). LL37 5 and 50 µg/ml (dark green and dark purple). N=3. (G) Survival of *S. aureus* after incubation with neutrophils primed with PBS, GM-CSF, HNP1, ClY21-HNP1, or IY16-HNP1 (all at 50 µg/ml). Data are derived from 11 healthy donors (N=11). Statistical analysis was performed using one-way ANOVA followed by Dunnett’s multiple comparisons test. (H) Representative live cell images of HMDM pretreated with HNP1, ClY21-HNP1 or IY16-HNP1 and then incubated with *C. albicans*^RFP+^ at MOI=5:1. HMDM were pre-stained with DAPI to label their nuclei. N=3. Scale bar, 50 µm.

**Figure S7.**
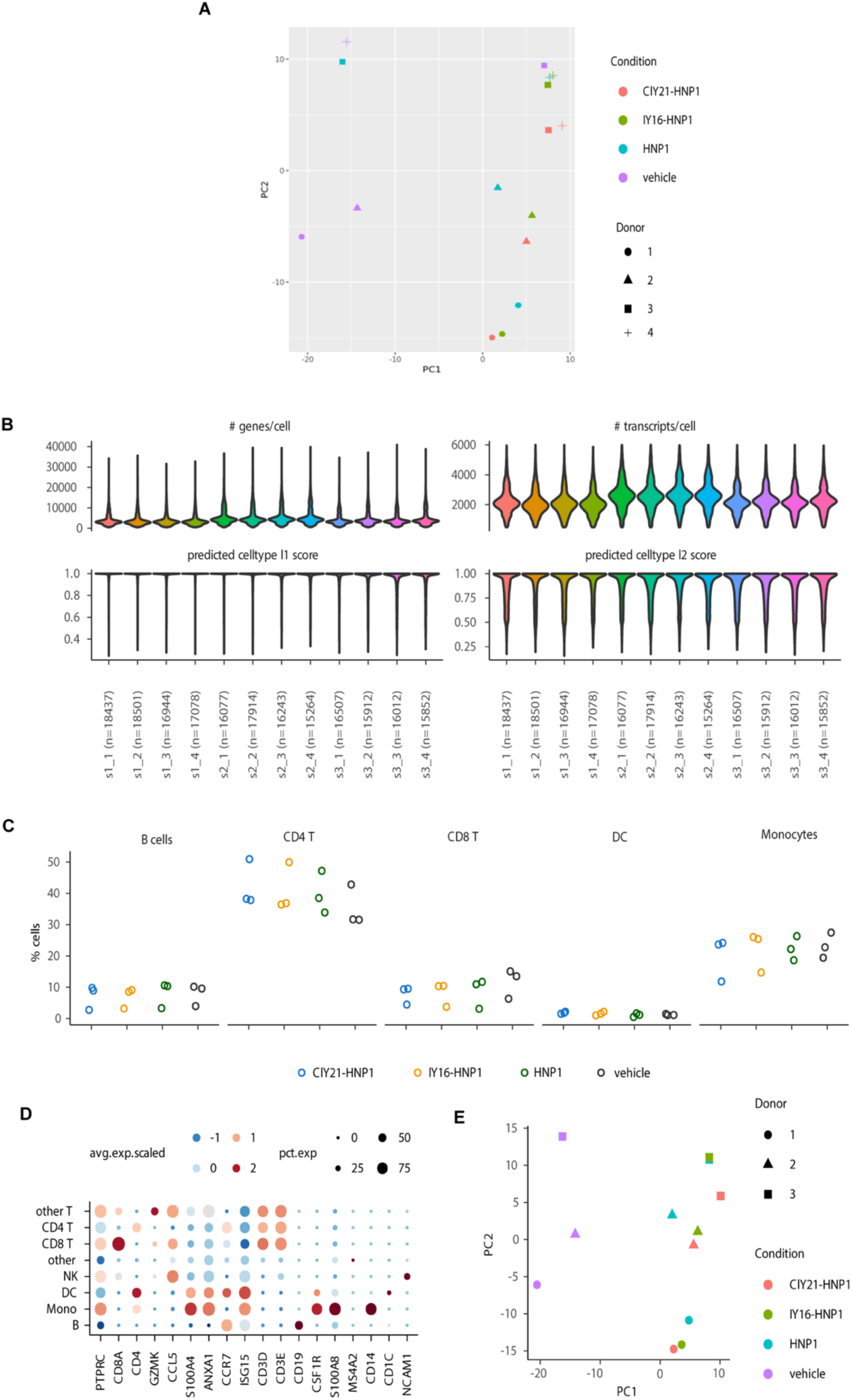
Halo-HNP1s are potent proinflammatory agents. (A) Principal component analysis of aggregated read counts from all cells (pseudobulk analysis) showing the principal components 1 and 2, where points are colored by the experimental condition group and shaped according to the donor. Donor 3 was excluded from the analysis because the data were identified as outliers, likely resulting from a sample swap affecting two of the four experimental conditions during processing. (B) Quality control of the processing and annotation of the scRNA-seq dataset using the human PBMC atlas reference mapping. For each sample, the distribution of number of detected transcripts per cell, number of detected genes per cell, as well as the label transfer scores for both layers of granularity (l1 and l2) are shown. The number of cells per sample is indicated with x axis labels. The labels indicate donor and condition e.g. d1_1 corresponds to donor1 and condition1. (C) Cell type percentages were calculated for each sample, in the four experimental groups (vehicle, ClY21-HNP1, IY21-HNP1, HNP1). (D) Dot plot showing the scaled average mRNA expression of selected marker mRNAs for the annotated cell types at the granularity level 1. Size of the dots correspond to the percentage of cells where marker mRNA was detected. (E) Principal component analysis of read counts from all cells (pseudobulk analysis) showing the principal components 1 and 2. Data points are colored according to the experimental group, while their shape corresponds to the donor.

**Figure S8.**
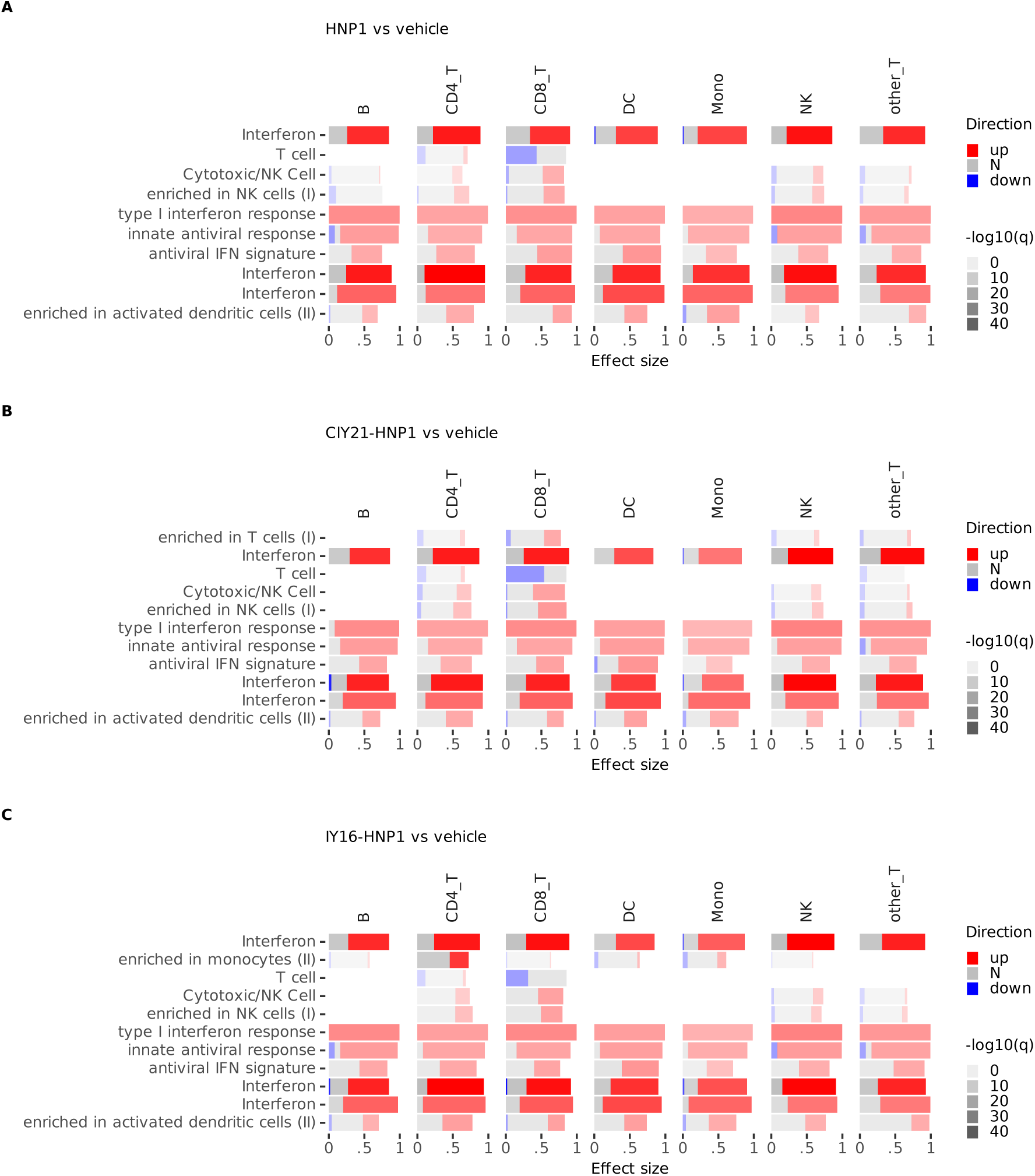
HNPs upregulate proinflammatory genes. (A-C) Gene set enrichment analysis of differentially expressed genes in indicated cell types (B cells, CD4+ and CD8+ T cells, dendritic cells (DC), monocytes, natural killer (NK) cells and other T cells). The comparisons are indicated above the panels and included HNP vs. vehicle (A), ClY21-HNP1 vs. vehicle (B) and IY16-HNP1 vs. vehicle (C). Bars represent relative fractions within each of the gene sets for mRNAs with decreased (blue) expression, non-differential (N) genes and increased (red) expression (determined by the adj. P value threshold of 0.05). Adjusted P values (q) for the gene set enrichment analyses correspond to the intensity of the bars, i.e. the lower the P value, the higher the bar intensity. Gene sets with AUC > 0.70 and q < 1e-10 are shown.

**Figure S9.**
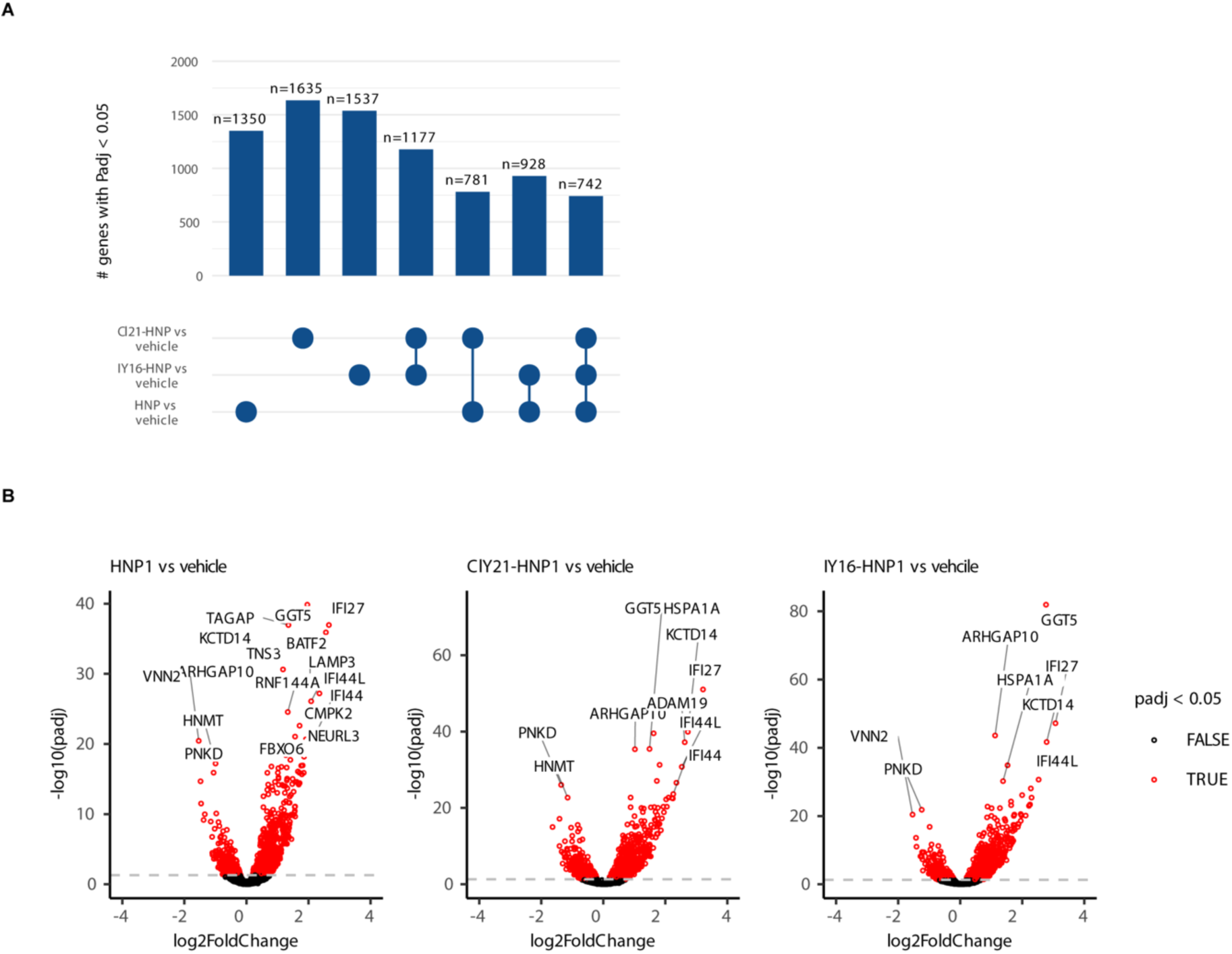
HNP1 and halo-HNP1s induce specific genes upregulation. (A) Upset plot depicting the overlap of differentially expressed genes (adj. P value < 0.05) between comparisons of different conditions. Absolute numbers are indicated above the bars. (B) Log2-transformed fold changes from the pseudobulk differential expression analyses for the HNP1, ClY21-HNP1 and IY16-HNP1 vs vehicle comparison in monocytes plotted versus negative log10-transformed adjusted P values (Wald test). Differentially expressed genes (adjusted P value < 0.05) are marked in red, while the non-differential (adjusted P value > 0.05) are shown in black.

**Figure S10.**
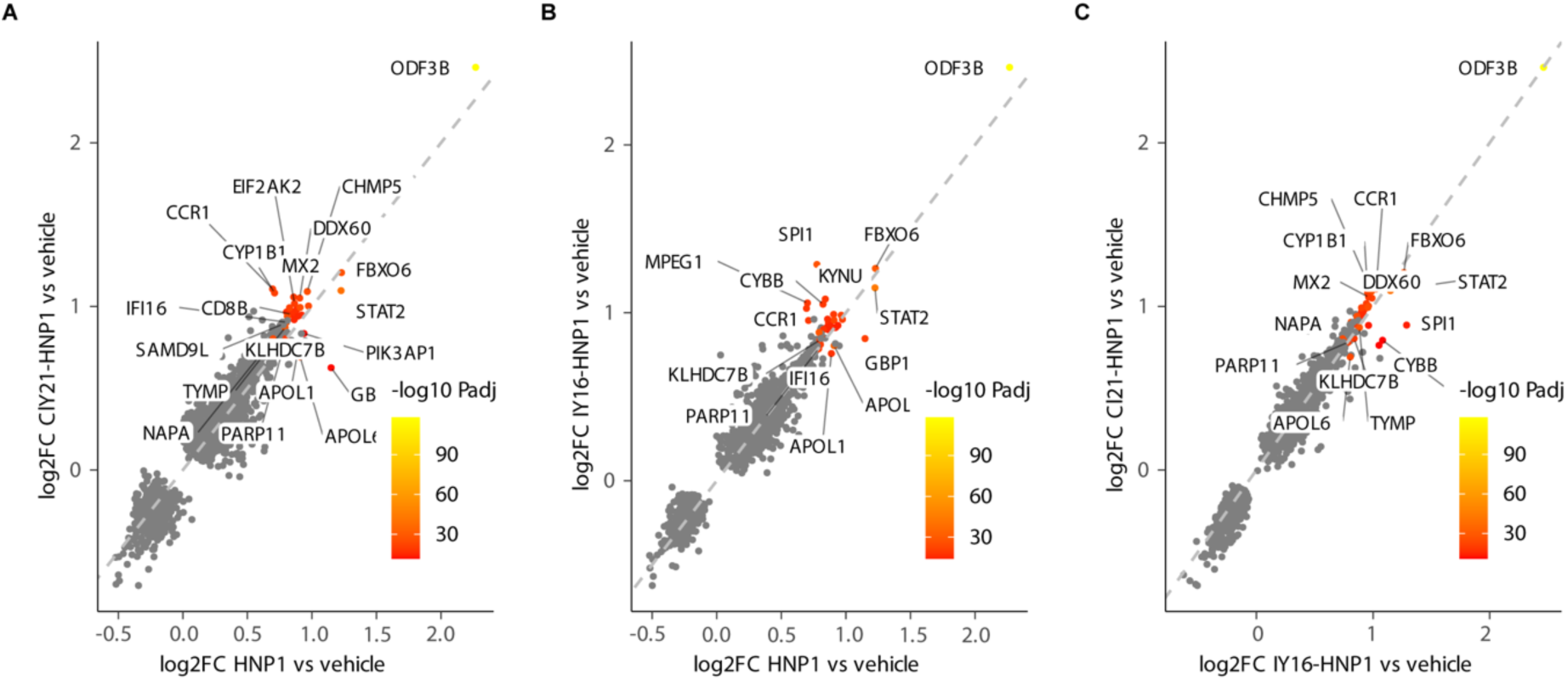
Halo-HNP1s modulate chemokine genes expression. (A-C) Scatter plots of log2-transformed fold changes (log2FC) derived from pseudobulk differential expression analyses in CD4 T cells. Dots represent all differentially expressed genes in either of the two depicted comparisons (Wald test, adj. P value < 0.05). Top 30 genes ranked according to discordance/concordance scores are depicted by colour gradient which corresponds to negative log10-transformed adj. P values in ClY21-HNP1 vs. vehicle (A), IY16-HNP1 vs. vehicle (B), or in ClY21-HNP1 vs. IY16-HNP1 contrast (C). Top 30 genes are also indicated with labels.

**Figure S11.**
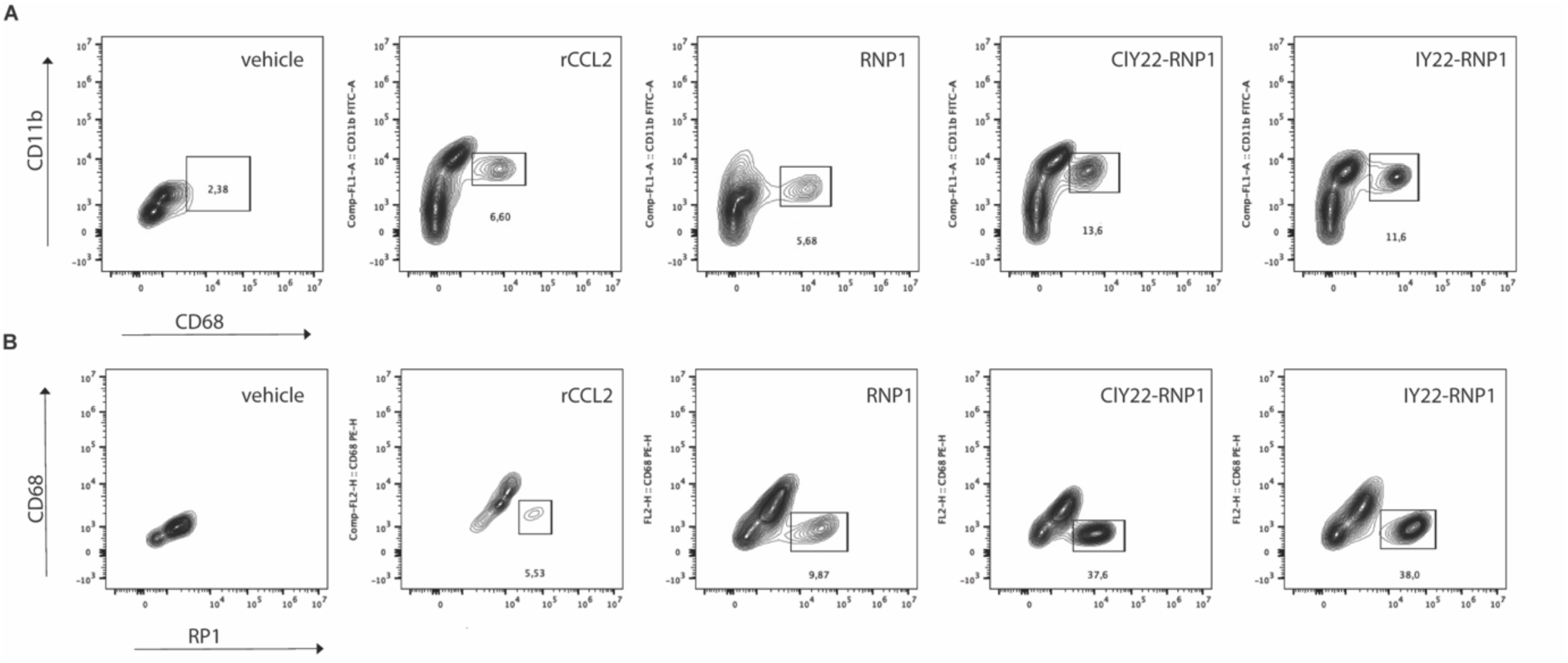
ClY22-RNP1 and IY22-RNP1 are potent proinflammatory agents *in vivo*. (A) Flow cytometry quantification of infiltrating monocytes (CD11b^high^ CD68+) 24 hour after injecting HNP1, ClY22-RNP1 and IY22-RNP1 or rCCL2, as a positive control, into the rat peritoneal cavity. Representative data of 5 independent experiments with 5 animals per group as shown in Figure 5 I. Statistical significance was analyzed in figure 5 I. (B) Flow cytometry quantification of infiltrating granulocyte (RP1^+^) 24 hour after injection of ClY22-RNP1 and IY22-RNP1 or rCCL2, as a positive control, into the rat peritoneal cavity. Data shown are representative of 5 independent experiments with 5 animals per group shown in Figure 5 J. Statistical significance was analyzed in figure 5 J.

**Table S1.**
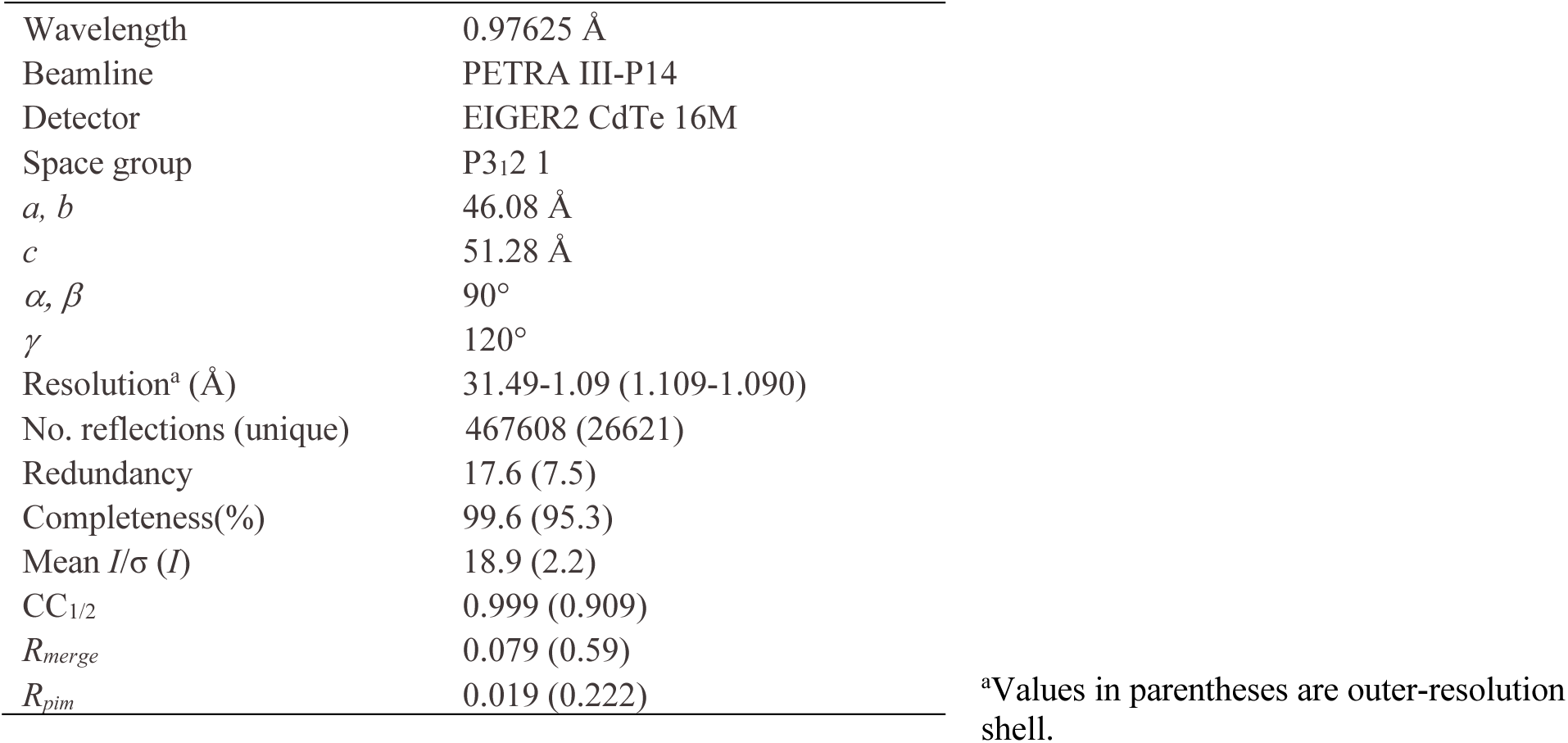
X-ray data collection statistics Cl21Y-HNP1.

|  |  |
| --- | --- |
| Wavelength | 0.97625 Å |
| Beamline | PETRA III-P14 |
| Detector | EIGER2 CdTe 16M |
| Space group | P3<sub>1</sub>2 1 |
| *a*, *b* | 46.08 Å |
| *c* | 51.28 Å |
| $\alpha$, $\beta$ | 90° |
| $\gamma$ | 120° |
| Resolution<sup>a</sup> (Å) | 31.49-1.09 (1.109-1.090) |
| No. reflections (unique) | 467608 (26621) |
| Redundancy | 17.6 (7.5) |
| Completeness(%) | 99.6 (95.3) |
| Mean *I*/σ (*I*) | 18.9 (2.2) |
| CC<sub>1/2</sub> | 0.999 (0.909) |
| *R*<sub>merge</sub> | 0.079 (0.59) |
| *R*<sub>pim</sub> | 0.019 (0.222) |

<sup>a</sup>Values in parentheses are outer-resolution shell.

**Table S2.** X-ray structure refinement statistics Cl21Y-HNP1.

|  |  |
| --- | --- |
| *R*-factor | 11.10% |
| *R*<sub>free</sub><sup>a</sup> | 12.18% |
| Solvent | 45.95% |
| Mean B-value (Å<sup>2</sup>) |  |
| Protein (main/side) | 23.67/28.08 |
| citrate | 21.64 |
| sodium | 33.73 |
| water | 36.43 |
| No. of protein residues | 1029 |
| No. of water residues | 740 |
| No. of citrate molecules | 8 |
| No. of sodium atoms | 1 |
| Root mean square deviations from<br>ideal geometry |  |
| Bond lengths | 0.015 Å |
| Bond angles | 1.595° |
| Ramachandran plot (%) |  |
| Favoured | 96 |
| Allowed | 4 |
| Outliers | 0 |

<sup>a</sup>*R*<sub>free</sub> was determined using 4.96 % of the data (71)

